# A Set of FMRI Quality Control Tools in AFNI: Systematic, in-depth and interactive QC with afni_proc.py and more

**DOI:** 10.1101/2024.03.27.586976

**Authors:** Paul A. Taylor, Daniel R. Glen, Gang Chen, Robert W. Cox, Taylor Hanayik, Chris Rorden, Dylan M. Nielson, Justin K. Rajendra, Richard C. Reynolds

## Abstract

Quality control (QC) assessment is a vital part of FMRI processing and analysis, and a typically under-discussed aspect of reproducibility. This includes checking datasets at their very earliest stages (acquisition and conversion) through their processing steps (e.g., alignment and motion correction) to regression modeling (correct stimuli, no collinearity, valid fits, enough degrees of freedom, etc.) for each subject. There are a wide variety of features to verify throughout any single subject processing pipeline, both quantitatively and qualitatively. We present several FMRI preprocessing QC features available in the AFNI toolbox, many of which are automatically generated by the pipeline-creation tool, *afni_proc.py*. These items include: a modular HTML document that covers full single subject processing from the raw data through statistical modeling; several review scripts in the results directory of processed data; and command line tools for identifying subjects with one or more quantitative properties across a group (such as triaging warnings, making exclusion criteria or creating informational tables). The HTML itself contains several buttons that efficiently facilitate interactive investigations into the data, when deeper checks are needed beyond the systematic images. The pages are linkable, so that users can evaluate individual items across a group, for increased sensitivity to differences (e.g., in alignment or regression modeling images). Finally, the QC document contains rating buttons for each "QC block", as well as comment fields for each, to facilitate both saving and sharing the evaluations. This increases the specificity of QC, as well as its shareability, as these files can be shared with others and potentially uploaded into repositories, promoting transparency and open science. We describe the features and applications of these QC tools for FMRI.

## INTRODUCTION

An important part of any FMRI processing pipeline is quality control (QC). QC has many aspects, which include:

1. evaluating the originally gathered data (e.g., for whole brain coverage, notable dropout, reconstruction errors or ghosting artifacts);
2. verifying that each step of processing completed successfully (e.g., adequate skull stripping, accurate alignments of volumes, reasonable statistical modeling);
3. reducing or eliminating errors in processing (e.g., incorrect timing files or units on timing, mistaken header information, blurring data with 40mm instead of 4mm radius);
4. and, generally, ensuring the validity and suitability of the acquired data for the subsequent analyses and interpretation.

Thus, while in many neuroimaging descriptions QC is simply referred to as the process of selecting out "good" or "bad" datasets to use in a group analysis, in the present work we view it as informing the researcher about the data properties in a fundamental way, as well as about the presence of possible artifacts or mis-steps in an analysis step. It should be used to assure the researchers that they fully understand the data at all stages of acquisition and processing, imbuing confidence that the subsequent analyses can be used to draw meaningful interpretations.

Increasing the size of data collections has the ability to increase statistical power and generalizability of results, and machine learning methods thrive on having large numbers of subjects as input. Many funding agencies also encourage data sharing, and repositories like the Functional Connectome Project (Biswal et al., 2010), UK Biobank (Sudlow et al., 2015) and OpenNeuro (Markiewicz et al., 2021) allow researchers to aggregate datasets from many sites and studies. As such, FMRI studies are increasingly including larger numbers of subjects, and the question arises about how best to evaluate the data quality and the processing results, balancing practical considerations with careful science. This is particularly challenging given that the FMRI blood oxygenation level dependent (BOLD) signal is known to be quite noisy and susceptible to sources of errors including various distortions, scanner artifacts, and more.

Ideally, every dataset in a study would go through a systematic QC process at the single subject processing level. Anecdotally, in all of the many collective years of experience working in neuroimaging, the present authors have never been involved in a FMRI project where such single subject QC was not needed. This was recently re-verified when looking at the contributions to the FMRI Open QC Project: many participating analysis Teams excluded at least 25% of the subjects in each of the 8 sites’ open data collections (Taylor et al., 2023a).

There are several forms that QC can take and several points at which to implement it. Indeed, verifying data as soon as it has been acquired at the scanner is the best starting point, particularly to reduce the possibility of acquiring inherently problematic data, whether due to non-ideal scanner settings, an underlying coil artifact, motion-inducing study paradigm or other issues. Here we focus on tools in the publicly available, open source software package AFNI (Analysis of Functional NeuroImages; Cox, 1996) for evaluating single subject data after initial processing, by which we mean checking data from its DICOM (the data format most commonly available from MRI scanner vendors) conversion through to individual statistical/regression modeling. Conveniently, this can also be combined with group-level quality assessment and evaluation, as we further describe. Rather than being separate from a processing pipeline, QC is itself an integral part of the processing within AFNI’s main FMRI analysis management tool, *afni_proc.py* (Reynolds et al., 2024).

From its initial design, *afni_proc.py*’s functionality has promoted the review of processing steps. Firstly, it creates a single subject processing script that is organized by processing blocks and commented—each step of processing is clearly laid out for description to others (including parameter values, such as blur radius, etc.) and for any later reference. When the processing script is executed, many intermediate data sets are created: these are not immediately deleted, but kept for the purpose of facilitating detailed review. In addition, QC scripts are automatically created for reviewing each single subject (ss) processing. For example, the "basic" review script provides a dictionary-like summary of informative quantities, such as maximum estimated motion, number of censored time points and TSNR (temporal signal to noise ratio); and the "driver" review script opens up an interactive guided review of key volumetric data in the AFNI GUI (checking alignment, statistical modeling results, etc.), as well as displaying plots of motion and regressors.

More recently, we have developed and expanded *afni_proc.py*’s QC (APQC) HTML, which is an automatically generated HTML document for systematically reviewing the FMRI processing steps for each subject. The APQC HTML incorporates and expands upon features of *afni_proc.py*’s "basic" and "driver" QC, containing volumetric images, both line- and box-plots of time series, and quantitative evaluations; from experience in FMRI analysis, we have found that both images and summary measures are necessary to understand a dataset. While a single scrollable document, the report is also organized thematically into "QC blocks" for assessing major features and steps of processing. The aim of this arrangement is to simplify recording and reporting of the QC itself. The APQC HTML was shown to be helpful in practice in evaluating data quality within the FMRI Open QC Project (see Reynolds et al., 2023; Teves et al., 2023; Lepping et al. 2023; Birn, 2023).

Here, we describe in detail the various layers and features of FMRI quality control tools in AFNI and how they can be applied in various roles of understanding the data and evaluating its suitability for analysis. We present several updates and new functionalities, particularly within the APQC HTML. These have been developed to increase both the efficiency and depth of data quality examinations, as well as the portability and shareability of the QC itself. For example, we have added a program to run the APQC HTML visualizations through a local server, so that QC ratings and comments/notes can be saved within the pages themselves in real time. Additionally, the webpage now contains buttons to open up the datasets quickly and efficiently for deeper investigation, both in the AFNI GUI and in a new embedded NiiVue panel (Hanayik et al., 2023). Automatic checks now include ROI-specific TSNR and shape properties, from both a default set of regions per template as well as lists of those the user can provide. Finally, we show how these qualitative evaluations can be combined with quantitative ones, facilitating systematic evaluations of data collections, using *gen_ss_review_table.py* and the new *gtkyd_check*. These features and others are demonstrated on real datasets, and we discuss these features in the context of QC within the FMRI field.

## METHODS AND RESULTS

As noted above, the primary goal of quality control (QC) in AFNI is to fully understand both the data’s properties from the beginning and its downstream results from processing (see Reynolds et al., 2024 for details). The standard *afni_proc.py* output contains features for different methods of investigation for a single subject’s (ss) data, ranging from individual automated checks and quantitative outputs to systematic snapshot to user-interactive views of the full datasets. This multilayered approach for evaluation allows the user to tailor their QC needs to a given collection of data, such as starting with fast quantitative checks and systematic image views, while also being able to efficiently dig more deeply into the datasets themselves to address any questions or concerns that arise. A brief summary of the primary tools described here is provided in Table 1. We describe these layers and features along with examples, using real FMRI datasets.

**Table 1.**
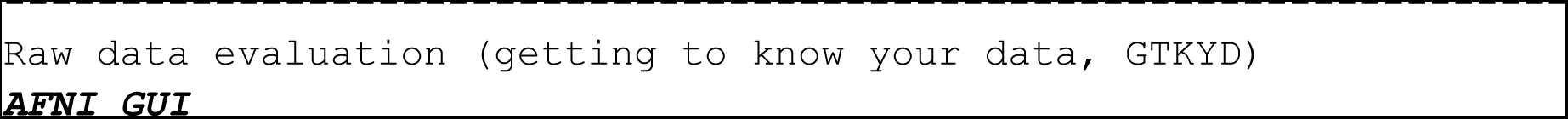
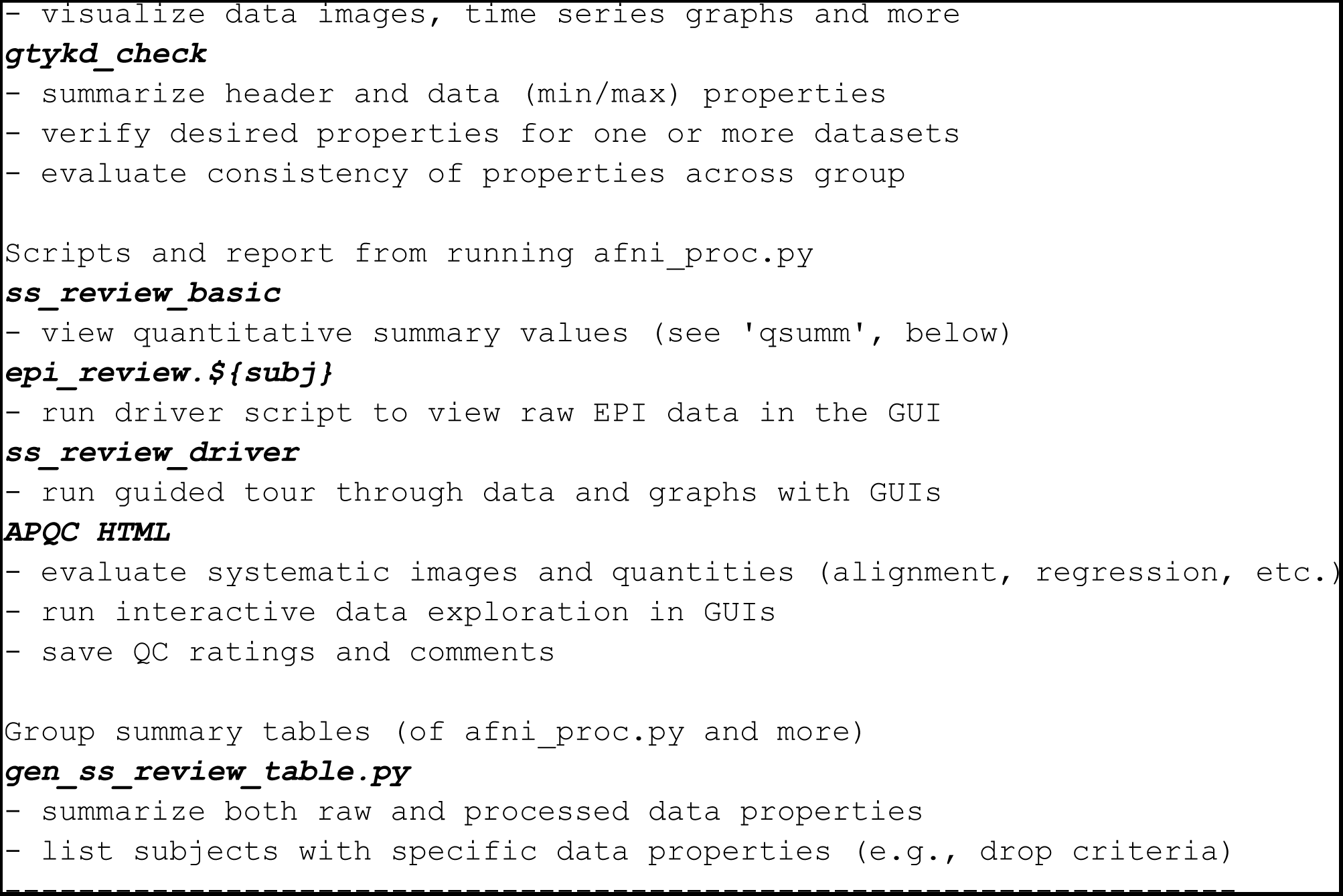
A brief outline and overview of the quality control functionality in AFNI discussed here.

The primary dataset visualized in the following examples comes from the basic AFNI "Bootcamp" class demo example (with original subject ID of two random characters "FT" there, but renamed to "sub-002" here), processed using AFNI v24.0.14. This task-based FMRI dataset has a block design for two timing classes: "vis", which is a visual-reliable stimulus (clear image with unclear audio), and "aud", which is an audio-reliable stimulus (clear audio with unclear image). This was processed using the newly available sswarper2 for nonlinear template alignment (Taylor et al., 2024) and a standard *afni_proc.py* script (see the Bootcamp’s afni_handouts documentation for details). Both the data and processing scripts are publicly available.^1^ For some QC examples, other publicly available data are also utilized, as described below.

### afni_proc.py output, I: processing provenance and intermediate datasets

To be able to evaluate the results of each processing block within the pipeline, *afni_proc.py* creates a standardized results directory that contains many intermediate datasets, as well as the copies of the original inputs and the final outputs. Intermediate datasets include results of various alignment steps, mask estimation, outlier estimation and censor files. Some further derived datasets of useful properties are generally estimated, such as temporal signal to noise ratio (TSNR), various correlation maps, and more.

Knowing the detailed provenance of the pipeline is important for understanding and accurately describing the processing. First, users are able to choose and specify a large number of options in processing; thus, seeing the details of the *afni_proc.py* command is the first level of understanding the processing in a relatively compact way; this is also publishable and shareable, facilitating reproducibility efforts (as long as the code version number is also specified, the processing pipeline can be fully recreated). Second, *afni_proc.py* does not simply process data and produce results, but instead it creates a full, commented script of all commands used so that the user can see exactly what happens at each step. Third, when using the "-execute" option, a log file of the terminal text is saved, to record the results of each command. Finally, each dataset in the results directory also contains the accumulated history of AFNI commands that have been used in its creation. In total, there is full provenance of the processing.

A JSON file "out.ss_review_uvars.json" is also output, which contains a standardized dictionary whose values are the names of important identification of output files and specific measures in the results directory; the keys define what each output is. For example, the "errts_dset" contains the regression model’s residual dataset (also known as the error time series), and the "sum_ideal" is a column file of the time series that represents the idealized response to all stimuli across the processed time series, for task-based FMRI data. The list of all possible keys, with brief descriptions of each, can be viewed with "gen_ss_review_scripts.py -show_uvar_dict". A sample subset of this is shown in Table 2, with the values of each key being set to values relevant for what was processed (code version number, names of output datasets through various stages of processing, etc.). This JSON file can serve as a guide for the user to identify important outputs, and provides a useful conceptual complement to other provenance features. It also is used by the AFNI programs that generate the HTML report for the processing, as well as other QC features. Being machine readable, this JSON file can be used in other/custom programs as well.

**Table 2.** A sample of the standard keys (uvar = "user variables"; first column) of important files and processing notes for the afni_proc.py results directory; the values are set by what was actually used/recorded in the processing (e.g., the AFNI package and version, the name of the anatomical dataset that was aligned to the EPI, etc.). Any of these may be contained within the results directory’s "out.ss_review_uvars.json" dictionary file, with its value providing the name of the relevant dataset or information. At present there are more than 50 keys in total (for full list, see "gen_ss_review_scripts.py - show_uvar_dict").

| **JSON (uvar) keys** | **: Brief description** |
| --- | --- |
| afni_package | : set AFNI package |
| afni_ver | : set AFNI version |
| align_anat | : anat aligned with orig EPI |
| censor_dset | : set motion_censor file |
| combine_method | : set method for combining multiple echoes |
| copy_anat | : original -copy_anat dataset |
| cormat_warn_dset | : correlation warns in Xmat |
| decon_err_dset | : 3dDeconvolve warnings |
| df_info_dset | : degree of freedom info |
| dir_suma_spec | : directory containing surface spec file |
| enorm_dset | : set motion_enorm file |
| errts_dset | : set residual dataset |
| final_anat | : anat aligned with stats dataset |
| final_epi_dset | : set final EPI base dataset |
| final_view | : set final view of data (orig/tlrc) |
| flip_check_dset | : -check_flip result dset |
| flip_guess | : guessed dset flip status |
| gcor_dset | : set gcor_dset file |
| have_radcor_dirs | : are there any (yes/no) |

### afni_proc.py output, II: automated quantities, checks and review scripts

The processing script created by *afni_proc.py* contains several places where automated checks for potential problems are calculated. These have been built up over many years of experience through helping users with troubleshooting various issues with data and modeling. Therefore, several logs of checks are kept within the results directory, including:

- left-right flip checks between the EPI and anatomical datasets (Glen et al., 2020)
- design matrix issues such as multicollinearity and ill-conditioning
- non-removal of pre-steady state time points in the EPI
- removal of too many degrees of freedom from the time series (e.g., due to subject motion, bandpassing or other processing choices)
- removal of too many time points from any stimulus class’s response interval (e.g., due to task-stimulated motion)
- through-plane variance lines, reflecting a scanner artifact.

Several of these checks combine quantitative evaluation with supplementary visualizations, to assist the user in validating the issue or understanding its extent more clearly. Each of these is also contained within the APQC HTML, and examples are provided below (Figs. 18-19).

A set of quantities of interest related to the subject data are also calculated by the *afni_proc.py* pipeline and stored within a text file in the results directory. This includes software information (operating system and AFNI version number), fundamental information about the input data (EPI TR, multiband level, slice timing pattern, voxel resolution, number of time points, etc.), processing-related information (number of censored volumes, degrees of freedom used, left-right flip evaluation, average motion, maximum motion, etc.), and useful derived quantities of interest (average TSNR within the brain mask, global correlation (GCOR; Saad et al., 2013), autocorrelation function (ACF) blur estimates (Cox et al., 2017), etc.). These are stored in a colon-separated file called^2^ "out.ss_review_${subj}.txt", with a subset of lines for an example shown in Table 3. This file essentially contains a list of potential quantities of interest to examine for absolute comparisons (is the TSNR too low? is the censor fraction of volumes too high?) or relative comparisons across subjects or groups; the method for tabulating and comparing these elements across subjects is described below.

**Table 3.**
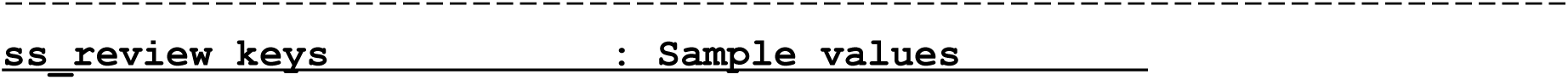
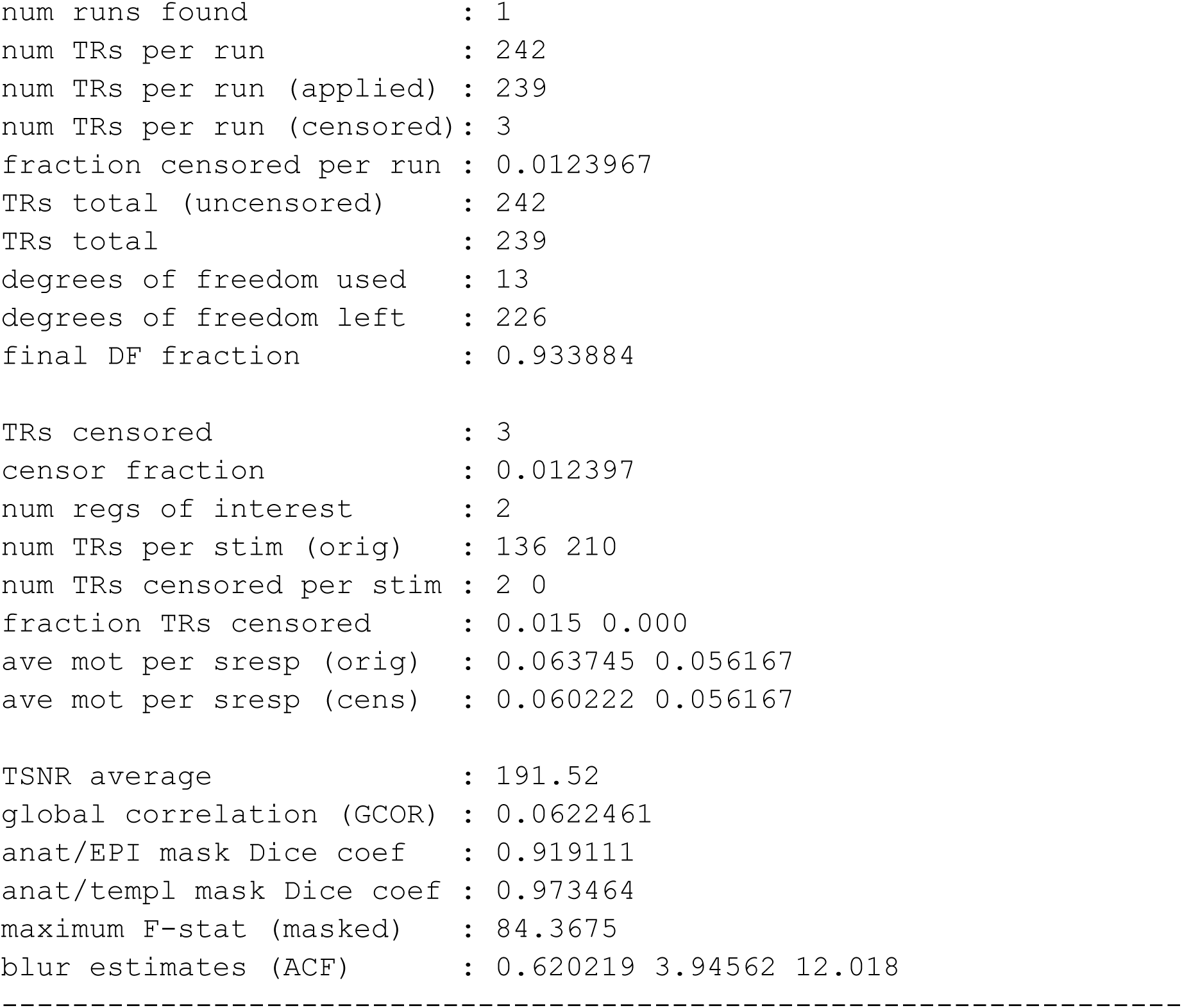
A partial sample of the standard keys (ss_review = "single subject review"; first column) of important quantities of input and processing-related information stored in the afni_proc.py results directory. These are contained within the results directory’s "out.ss_review_${subj}.txt" dictionary file, with its value providing the relevant quantity or information. At present there are more than 50 keys in total. Above, "sresp" quantities refer to each stimulus response (specifically, to time points where the idealized response regressors are nonzero).

Visualization plays a key role in understanding data, complementing quantitative checks, and *afni_proc.py* creates several scripts that facilitate interactively reviewing datasets, information and related plots. The first review script added (in 2008; *afni_proc.py* was created in 2006, see Appendix B of Reynolds et al. (2024) for a timeline of some feature additions) starts with *@epi_review.${subj}*, and it drives the AFNI GUI to show the user image and graph views of the input EPI datasets, automatically scrolling through time in video mode. Another set of review scripts (added in 2011) start with "ss_review" for single subject review. Running *@ss_review_basic* creates and displays the "out.ss_review_$ {subj}.txt" text file, described above, for quick review of basic dataset quantities. Running *@ss_review_driver* makes use of the "driveable" structure of the AFNI (and SUMA) GUIs. It guides the user through a set of plots and datasets to review, successively bringing up each plot and a descriptive text panel for each:

- Text display of the "basic" review quantities (described above)
- Plots of estimated subject motion, time series outlier fractions, and censoring
- Alignment of the anatomical and EPI datasets (viewable in the AFNI GUI)
- Text displays related to the general linear modeling: degree of freedom usage and potential regression warnings
- Plots of the individual and summed ideal stimulus responses (for task-based FMRI)
- Overlays of statistical results in the AFNI GUI; the volume of complete statistical results is loaded, and the full F-stat is overlaid, for evaluating the omnibus fit measures; the user can change the overlay and threshold indices to display both the effect estimate (beta coefficient) and statistical information, respectively (see Chen et al. (2017) for the importance of using both) of any additional stimuli or contrasts.

Some of these are shown in Fig. 1. Finally, running *@ss_review_driver_commands* contains the same driver commands, but more compactly, for a user to copy-and-paste or generally learn from. In this way, the user can quickly evaluate many of the key aspects of the processing pipeline, while not being constrained to a subset of static slices, colorbar/range choices, etc.

**Figure 1.**
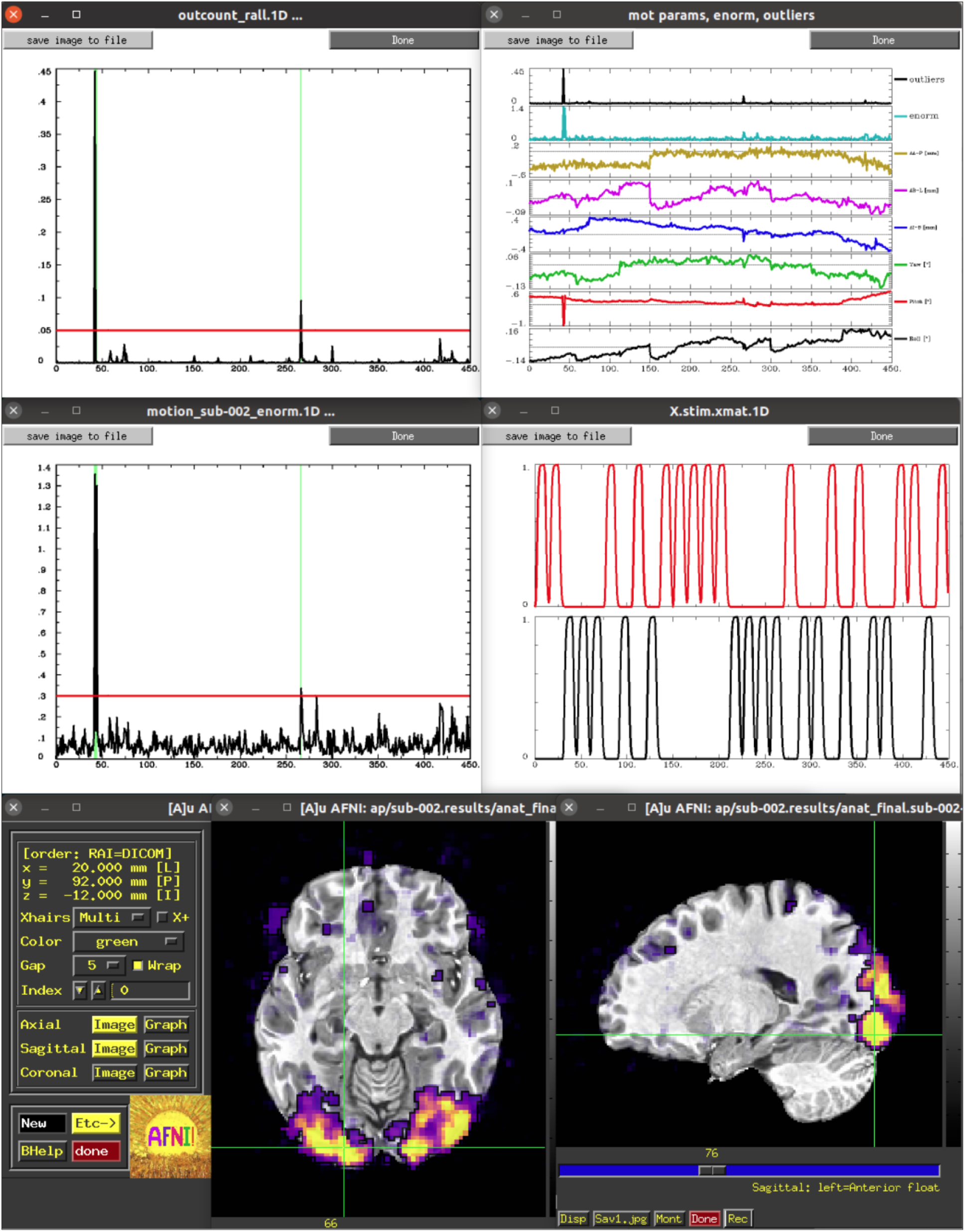
A selection of the plots and GUI views of single subject data from the ss_review_driver script. These include: (top) time series plots of Enorm (the scalar norm of estimated subject motion) with censoring and the individual motion regressors (with Enorm and outlier fraction); (middle) time series plots of the fraction of outliers within the brain volume (with censoring) and of the idealized stimulus responses for the two block-design classes in the FMRI study; (bottom) an interactive AFNI GUI view of the general linear model statistics, starting with the full F-stat and using transparent thresholding (Allen et al., 2012; Taylor et al., 2023a).

More recently, a set of scripts that start with "run_qc" have been added to the results directory output, as well. Each of these drives the AFNI GUI to visualize a particular processing aspect, displaying a montage instead of a single-slice view in each plane. The list of features viewed is an expanded set of those in the *ss_review\** scripts, including TSNR, different correlation maps, and per-effect visualizations of statistical information (so the user does not even have to change indices to select different stimuli or contrasts of interest). These may be run directly via the command line by the user, but they add particularly convenient utility to the APQC HTML, as described below.

### afni_proc.py output, III: interactive APQC HTML

One of the last outputs produced by the pipeline script is *afni_proc.py*’s QC report, which is referred to as the APQC HTML. This is created in the results folder in the directory "QC_${subj}". This directory contains the systematic quality control information for review. The APQC HTML can be viewed using any reasonably modern web browser (no internet connection needed). The APQC directory is typically 10-12 MB in size and portable: it can be moved and viewed on other computers, sent to collaborators or made available online with publicly shared results.

The APQC report is a single HTML page ("index.html") in the QC directory that contains a wide assortment of systematic information, with the goal to comprehensively understand the data from its raw state through full subject-level processing, which in AFNI is typically the linear regression modeling from *afni_proc.py’s* "regress" block. The page is organized as a sequence of major "QC blocks," each of which typically contains multiple items, which are some combination of images and/or text. The APQC HTML also contains a link to a help document within the QC folder for easy reference, and it can also have links to subpages of QC documents from other software that have been called via *afni_proc.py*. It also contains interactive functionality for the user to both directly open up datasets in GUIs and save comments and QC ratings, as described below. We have created an online demo of the APQC HTML (https://afni.github.io/qc-demo-repo/), so that users can directly explore some of the systematic images and interactive functionality.

At the top of the HTML page is a permanently visible menu bar (Fig. 2), which contains several buttons and text items. Hovering over each button provides a brief description. The top row provides a label for each QC block within the page, on which the user can click to jump to that block in the page (the HTML page contents for each QC block are detailed below). Beneath each label is a button, with which a user can save two types of QC evaluation information: a (quick) rating and a (detailed) comment. As described in Fig. 2B, the following properties apply to the QC rating and comment buttons:

- To save a rating, the user simply clicks on the button to toggle through a list of evaluations: good (+), bad (X), and other (?). For speed, there are "filler" buttons for each rating category located at the right side of the menu: a single click fills *empty* rating buttons, and a double-click will fill *all* rating buttons (even overwriting values). The "FINAL" label at the top does not link to a particular QC block, but allows the user to store an overall evaluation of the QC for this subject.
- To save a comment, the user can Ctrl+click (on macOS, Cmd+click) on the button, which opens a text box panel. The user can simply type any desired notes or questions and hit "Enter" to immediately save the information (and close the panel). A pair of quotations will appear around the value of the rating for which a comment is saved, so that it is visually apparent that a comment has been saved. The panel can be similarly reopened to review, edit or remove the comment. When saving a comment to an unrated QC block, a rating of "other" will automatically be applied.
- Comments can be cleared via a button in the text field, or by hitting "Esc" while the cursor is in the text. The individual QC button information (any comment and/or rating) can be cleared via the third button in the text field, or by hitting "Ctrl+Esc" while the cursor is in the text.

**Figure 2.**
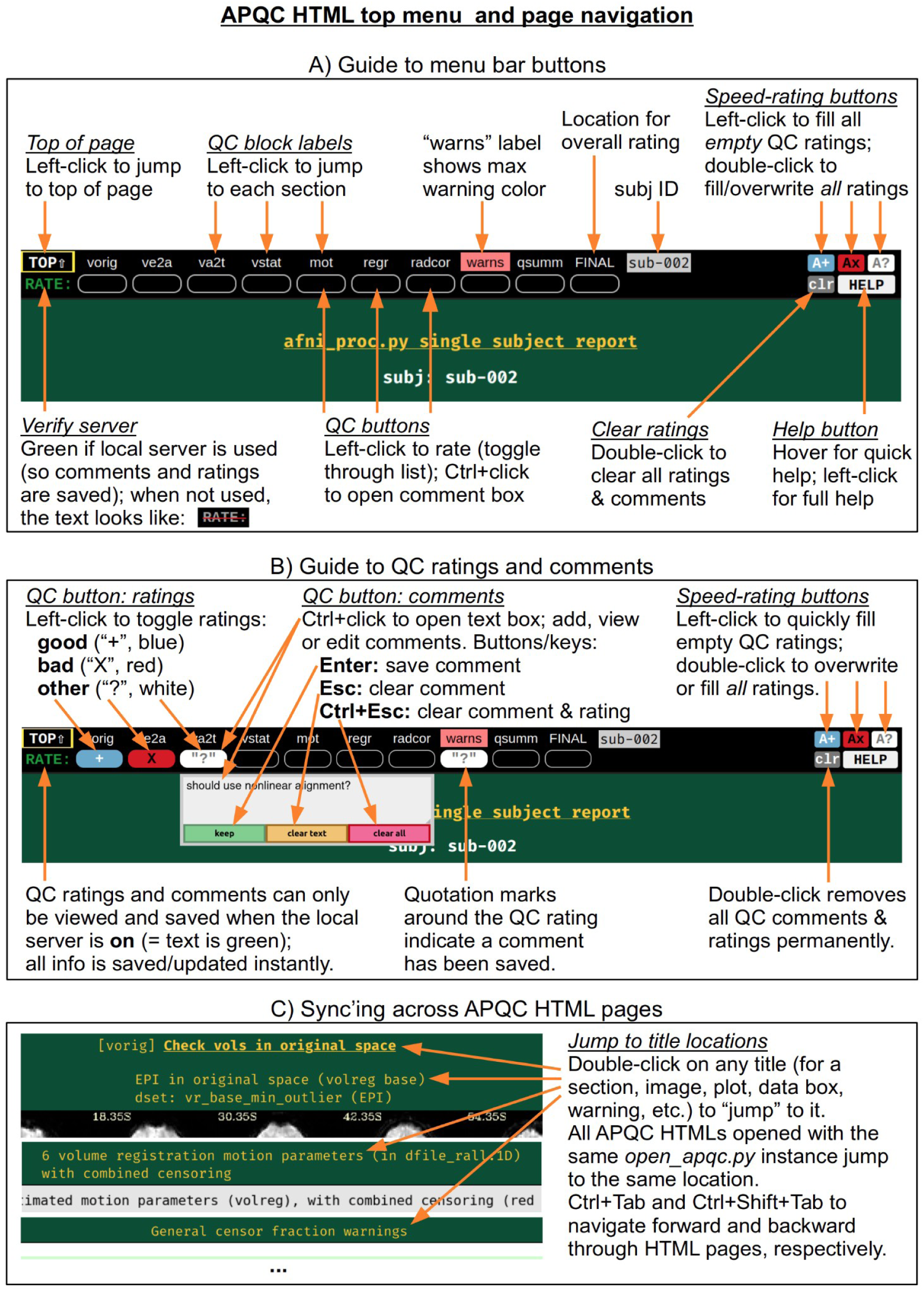
A quick guide to button features within the top menu bar of the APQC HTML. The menu bar is permanently visible on the page. The QC labels can be clicked on to jump to any QC block on the page. When the HTML is viewed with an active local server, the buttons beneath each label save and display QC ratings and descriptive comments. This functionality facilitates the storage and sharing of data evaluations. Jumping to the same location and syncing across multiple APQC HTML pages provides for highly efficient QC evaluation and comparison, particularly for images (alignment, effect estimate and statistic maps, etc.).

All rating and comment information is immediately saved in a JSON file in the QC directory. The comment functionality is particularly useful for storing free-form evaluative thoughts in real time, and allowing the user to share them later. We describe below how the rating information can be quickly parsed and compared across subjects, such as for listing all those with a given rating; this might be useful when implementing drop criteria, or when looking for patterns across subjects.

For speed, one can also navigate the top menu using the "Tab" key. The "Enter" and "Ctrl+Enter" keystrokes behave just as left-click and Ctrl-click, respectively, to make ratings and comments. There is a "Help" button at the right of the menu bar, providing references such as descriptions of QC blocks and abbreviations, as well as summaries of keystrokes and links to online documentation and resources.

The QC rating and comment information are stored in text files within the QC directory. Due to modern browser security, interacting with files on the local computer is greatly facilitated by running a local server. In AFNI, this is accomplished using the Python modules Flask and Flask-Cors, and the functionality works across Python versions 2.7 and 3.*. Once the modules are installed, the AFNI program *open_apqc.py* can be used to open one or more APQC HTML files, opening each in either a new tab or window of the default browser. The "RATE" text in the upper left shows that the pages were successfully opened by appearing with green font. When the pages have been opened in this way, the QC rating and comment buttons work and show any information saved from previous sessions.^3^

It is also possible to jump to a particular QC block or item immediately when opening the HTML. One provides the name of the appropriate location via "open_apqc.py -jump_to..". Each QC block label within the top menu is a possible jump location, and the full list for a page can be displayed with the "-disp_jump_ids" option. This is particularly useful for opening multiple APQC HTML pages at the same QC item in multiple tabs, and then browsing across them with standard "Ctrl+Tab" and "Ctrl+Shift+Tab" keypresses to navigate forward and backward through the tabs. One can also double-click on any title (gold text) within an HTML page to "jump" to it in both that page and all other APQC HTMLs that were opened with the same *open_apqc.py* instance (Fig. 2C). In this way, patterns of similarity and variability across the subjects is readily visible in a highly efficient manner, while still being able to save comments and ratings for each page individually.

We now describe the currently available set of QC blocks, which are listed with brief overviews in Table 4. Many of these contain images of volumetric views, which are typically shown as montages of evenly-spaced slices across the volume. A set of axial views is always shown, and in most cases sagittal images are also included. Each panel shows a full slice, but slices are selected from within a light automasking of the FOV (field of view); this reduces the chance of "wasting" a slice view on empty FOV. The slice coordinates are labeled. The filename of the volume(s) in the image is also provided, along with a contextual description, so that the researcher can view it more in-depth in the results directory, if need be. In general, these images provide a systematic view of a wide variety of features in the data, from its raw origins through intermediate steps and final stages for the single-subject level of processing. Because they are systematic and multi-slice, they provide an efficient way to visually check features within a subject and to compare across subjects.

**Table 4.** A brief outline and overview of the QC blocks in afni_proc.py’s APQC HTML. More blocks and items will likely be added over time. For a more complete list of QC items to consider evaluating within each block, see the main text here as well as Reynolds et al. (2023), Birn (2023), Lepping et al. (2023) and Teves et al. (2023).

| Block | Example features to evaluate |
| --- | --- |
| **vorig** | EPI and anatomical FOV, coverage, distortion, tissue contrast, brightness inhomogeneity, initial EPI-anatomical overlap |
| **ve2a** | EPI-anatomical alignment, EPI coverage, distortion |
| **va2t** | anatomical-template alignment |
| **vstat** | effect estimate + statistics maps, correlation maps, artifacts, expected/unexpected patterns (via InstaCorr, Graph View, etc.) |
| **mot** | subject motion, stimulus-correlated motion |
| **mecho** | multi-echo fMRI features and processing |
| **regr** | stimulus regressors (if present), corr_brain, TSNR patterns |
| **radcor** | local correlation patterns across the brain, InstaCorr of EPI at different stages |
| **warns** | pre-steady state, left-right flip, variance artifacts, high subject motion, regression correlation, etc. |
| **qsumm** | quantitative summary values (see gen_ss_review_table.py) |

For each volumetric image, there may be up to four interactive visualization buttons that can be used to go beyond the static image. When the QC directory is still located within the *afni_proc.py* results directory, clicking on of the buttons results in one or more datasets being loaded into either the AFNI GUI or a NiiVue instance (Hanayik et al., 2023) embedded within the browser, which can then be explored interactively. The behavior of the buttons are detailed in more detail below in the description of the "vstat" QC block and in Figs. 6-9, but briefly they are:

- **AV**: load the AFNI GUI viewer of the montages to navigate through
- **GV**: view time series in the AFNI Graph Viewer
- **IC**: surf seed-based correlations with AFNI’s InstaCorr (Song et al., 2017)
- **NV**: open NiiVue within the HTML to browse data.

### vorig: original EPI data

This block displays the original EPI and anatomical datasets in their original spaces (Fig. 3). Viewing the original data contains useful QC information: is the entire brain in the FOV (if intended)? Is there severe brightness inhomogeneity or distortion? Is the tissue contrast appropriate? Is the brain upside down (yes, this can occur in mainstream, public repositories; see Reynolds et al. (2023) for examples)? In addition to the grayscale colorbar labeling, ranges of minimum and maximum values are also presented: occasionally, negative values can be signs of a problem in data acquisition or conversion; additionally, having a maximal value of 4095 might be a warning of having saturated the 12-bit intensity signal (see e.g., Sakaie and Lowe, 2006) during acquisition.

**Figure 3.**
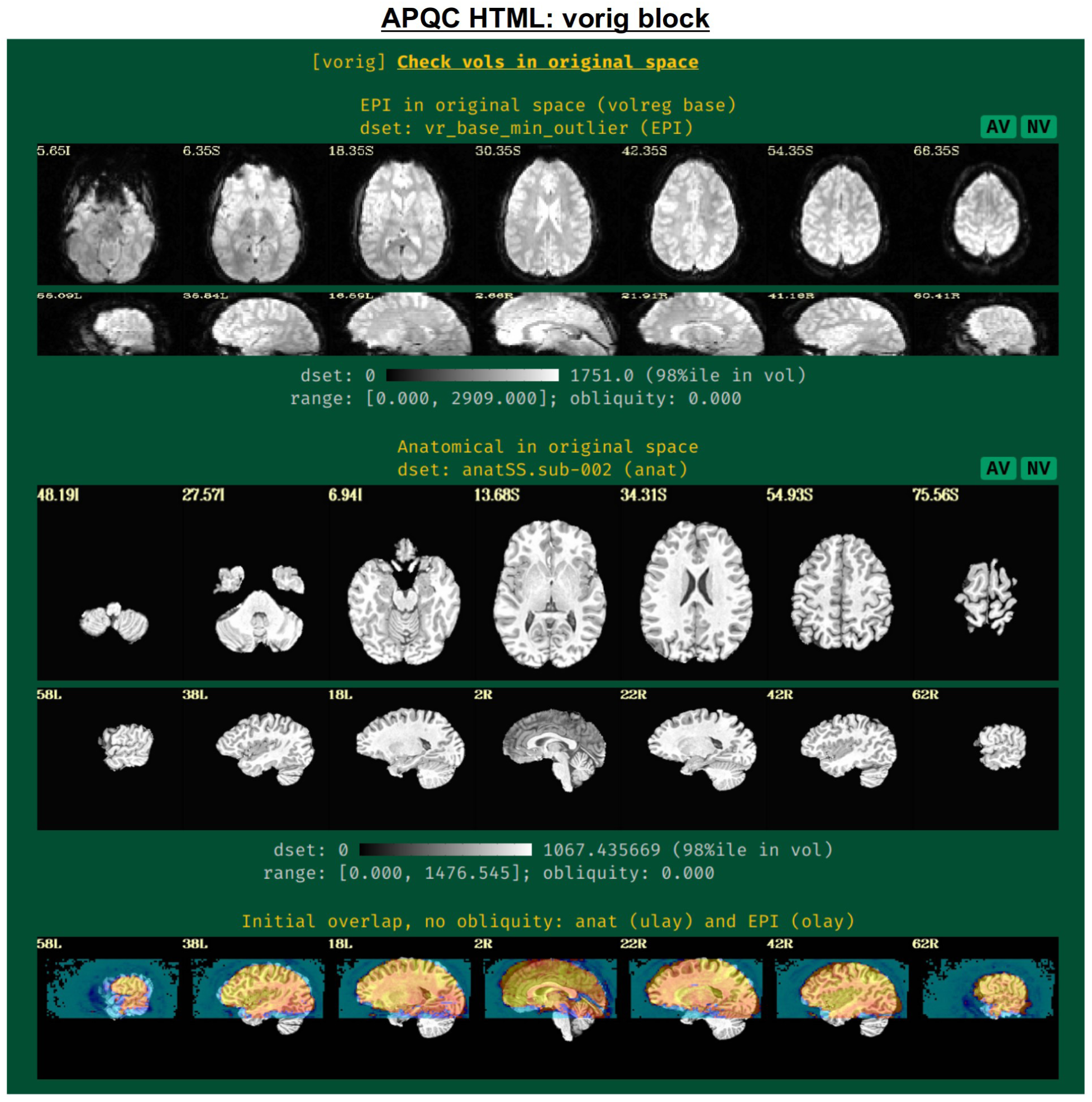
Example information within the "vorig" QC block. This displays an example volume of the raw input EPI and anatomical volumes, which can be assessed for FOV coverage, tissue contrast, distortion and other features. Seeing initial EPI-anatomical overlap (bottom row) can help troubleshoot their alignment during processing, if issues occur. The AV and NV buttons are part of the interactive APQC HTML functionality, in which the respective datasets can be opened in full GUIs if further examination is needed, described below in Figs. 6 and 9.

The displayed volume of the EPI time series here is typically the one that has been selected for both alignment to the anatomical volume and as a motion correction base. Reynolds et al. (2024) recommend using the AFNI functionality to choose this based on finding the one with minimal outliers across time (using the recommended "-volreg_align_to MIN_OUTLIER" in *afni_proc.py*). The quality of this determination can also be visually verified in this QC block.

An image of the overlap of the initial EPI and anatomical datasets in their native spaces is also shown. If EPI-anatomical alignment fails, this image may help troubleshoot the issue. If the datasets start "far" from each other, they are more likely to be stuck in a local minimum of poor alignment; a remedy could be to reset their coordinates, open up the parameter space of alignment, or check for DICOM conversion errors that produced erroneous coordinates.

### ve2a: alignment of subject EPI to subject anatomical

In nearly all processing, whether clinical or research, the EPI volumes are aligned to the subject’s anatomical scan (typically a T1-weighted image with full brain coverage and high spatial resolution in all dimensions). Due to the field-of-view, higher resolution and contrast of the anatomical, this allows for easier identification of structures within the EPI and a better basis for aligning it to template space, if necessary. Since alignment is performed using the best quantitative evaluation for the job (that is, the cost function), the quality of its results is best verified visually (Fig. 4, upper panel). Having "good alignment" is determined by having similar internal structures in the same location, such as ventricle edges, tissue boundaries, gyral/sulcal patterns, etc. Here, the edges of the higher resolution and more spatially detailed anatomical are shown on the EPI. This provides a clear picture of the relationship between most internal structures in the functional and anatomical images.

**Figure 4.**
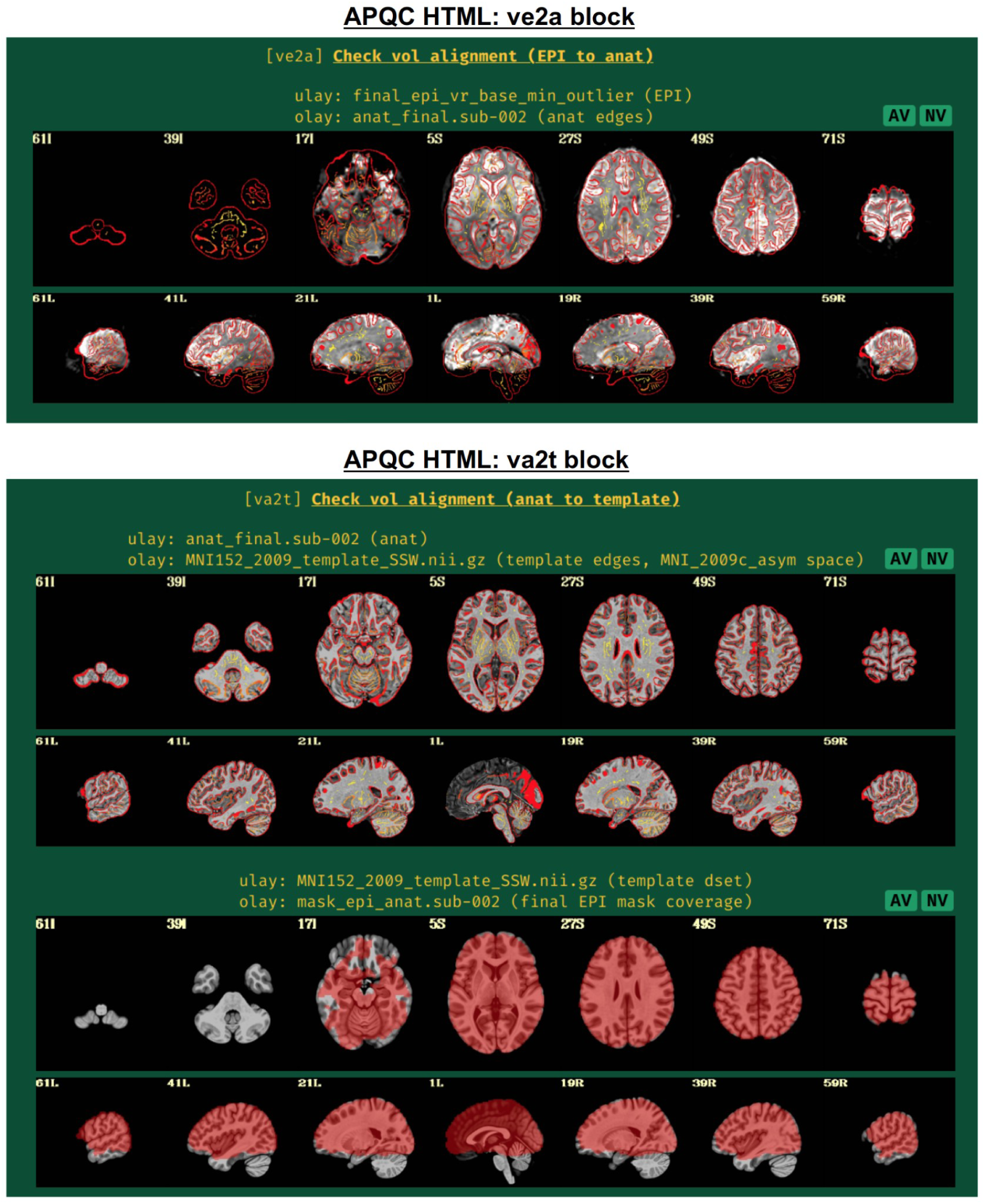
Example information within the "ve2a" and "va2t" QC blocks. These display results of intermediate processing, namely alignment of EPI-to-anatomical and anatomical-to-template datasets, respectively. Visual assessment is typically needed to provide final evaluation of the quality of the algorithm; here, edges of one dataset are shown over the other. Note that beyond alignment, the ve2a image provides useful information on EPI coverage, distortion and signal attenuation. The EPI coverage mask that afni_proc.py estimates during processing (to use in calculations, but not apply to hide EPI data) is shown in the bottom panel. It is helpful to see where EPI coverage is relatively strong or potentially missing. The AV and NV buttons are part of the interactive APQC HTML functionality, in which the respective datasets can be opened in full GUIs if further examination is needed, described below in Figs. 6 and 9.

Poor alignment could result from having low or non-standard tissue contrast, which may be resolvable by choosing a different cost function. Additionally, poor initial overlap can lead to false solutions of alignment (hence it is shown in the "vorig" QC block), which may be aided by fixing problematic coordinates in the file headers. AFNI alignment programs also have several options for incorporating a wider parameter space into the optimization process.

Additionally, this QC block can also show very useful information about signal coverage for the subject. For example, is the signal weak in the ventromedial prefrontal cortex and/or subcortex, which is very common in FMRI? It can also highlight potential local or global distortions, with the T1w edges providing useful anatomical reference information.

### va2t: alignment of subject anatomical to template

This QC block shows volumetric views of alignment between the subject anatomical and a template dataset, if selected (Fig. 4, lower panel). To facilitate comparison, the edges of the anatomical are overlaid on the template. As nonlinear registration is typically recommended for this step in AFNI’s FMRI processing (Taylor et al., 2018), the quality of alignment in this case should be quite high, whether using AFNI’s nonlinear warping tool *3dQwarp* (Cox and Glen, 2013) or one of the recommended wrapper programs that combine skullstripping (ss) with nonlinear alignment (warping), such as *@SSwarper* or sswarper2 for human datasets or *@animal_warper* for nonhuman datasets (Jung et al., 2021); each of these programs’ results are easily integrated into *afni_proc.py* (Taylor et al., 2018; Reynolds et al., 2024). Again, this alignment procedure requires visual verification of the algorithm’s results (and see the comments on potential alignment issues, in "ve2a", above).

If the "mask" block within *afni_proc.py* was used, a brainmask is calculated (Fig. 4, lower panel, bottom row). Typically, this is an intersection of the anatomical and the EPI data, locating where signal strength is relatively high. This mask is used for calculations of within-brain statistics such as whole-brain average TSNR, but importantly is not applied to create a separate masked-out EPI dataset; this latter procedure would lose valuable information for both data understanding and EPI quality evaluation, as well as hide potential artifacts (Taylor et al., 2023b).

### vstat: basic statistical modeling

The individual subject processing in *afni_proc.py* also includes regression modeling, whether task-based or resting/naturalistic. For task FMRI, 3dDeconvolve’s full F-stat, which is essentially the ratio of variance explained in the model’s regressors of interest to the variance of the residuals, provides a useful assessment for the quality of model selection (Fig. 5, top panel). In particular, one might look to see that its values are high in the regions of the brain which should be associated with a given task (e.g., in the visual cortex for flashing checkerboard task, etc.). Issues observed here may arise from having scanner artifacts in the EPI data, stimulus-correlated subject motion, poor task compliance by the subject, incorrect timing in the stimulus files, or other sources.

**Figure 5.**
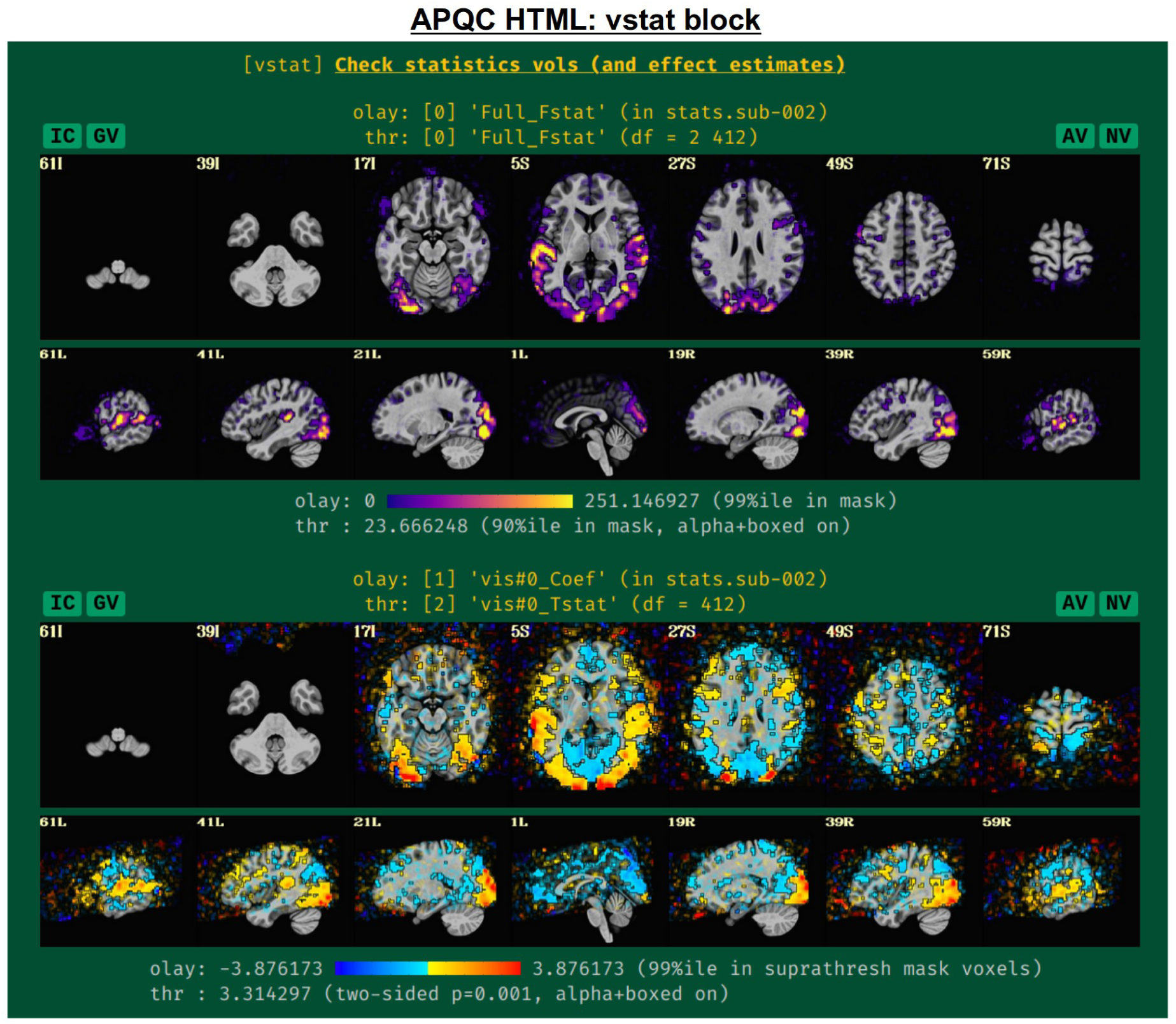
Example information within the "vstat" QC block. For task-based FMRI, the full F-stat of the single-subject regression modeling is shown first. Then, the effect estimate and statistical information for individual stimulus classes are shown, as well as similar maps for contrasts (general linear tests) the user had asked for; by default, the F-stat and up to 4 further statistical images are shown, but the user can add more. The datasets are not masked, and the "highlighting" methodology of thresholding is applied—values above threshold are opaque and outlined, while those below become increasingly transparent—to greatly enhance interpretation and understanding of the full datasets (Taylor et al., 2023b). The IC, GV, AV and NV buttons are part of the interactive APQC HTML functionality, in which the respective datasets can be opened in full GUIs if further examination is needed, described below in Figs. 6-9.

The range of the colorbar is shown below the image, along with the rationale for how it was selected automatically. The max value is the 99th percentile value in the whole brain (WB) mask, while the threshold is the 90th percentile value in the WB mask. An important note is that the threshold here is applied transparently to "highlight" suprathreshold voxels, while not hiding other features that might be interesting or alarming, and hence worth knowing about (Allen et al., 2012; Taylor et al., 2023b). This is done using AFNI’s "alpha" setting in the visualization: suprathreshold voxels are opaque, and subthreshold voxels have an opacity that decreases with their value (i.e., their "alpha" channel decreases quadratically from the threshold). To further highlight the suprathreshold voxels, these regions are bounded by a black line (i.e., “boxed”). Additionally, the overlay is not restricted to a brain mask, because it is often useful to see the results in the whole FOV: while there might be extraneous noise features outside the brain, it is useful to know if there might be ghosting, bad coil features, poor alignment, or some other pathology in the data. This form of visualization is particularly useful for QC and not hiding detrimental features unnecessarily.

Within this QC block, additional volumetric images are shown for the individual stimulus components, as well as specific general linear tests (GLTs) or contrasts of interest that the user might have included in the *afni_proc.py* command. For GLTs with both an effect estimate (beta or coefficient) and a t-statistic, the former is displayed in the overlay and the latter is used to threshold, following Chen et al. (2017); the 2-sided t-statistic (Chen et al. 2019) is thresholded at *p=*0.001, with the alpha+boxed properties described above. By default, up to 4 of these additional stimulus and GLT datasets will be shown in this QC block, in addition to the full F-stat volume, but the user can choose to add more when running *afni_proc.py* (via "-html_review_opts").

In Fig. 5, there are four buttons visible above the image montages: AV, GV, IC and NV. Most other images have these above them, as well. (If the HTML page is opened without the local server being active, these are not active, and instead will only be hinted at by appearing as empty green buttons slightly lighter than the background; moreover, the buttons can only connect to the datasets in question when the QC directory is inside its parent *afni_proc.py* results directory.) Each of these buttons allows the user to efficiently dig more deeply into the data, such as to browse through the full set of slices or to investigate different attributes. Brief descriptions and visualizations of these buttons’ functionality are shown in Figs. 6-9 and described in each caption. The AV, GV and IC buttons each makes use of several of the run_qc*tcsh scripts (mentioned above) that were created by *afni_proc.py* and which sit in that pipeline’s results directory. In short, viewing systematic images is useful for quick evaluations, there are many times when it is quite helpful to be able to quickly follow-up on any questionable features.

**Figure 6.**
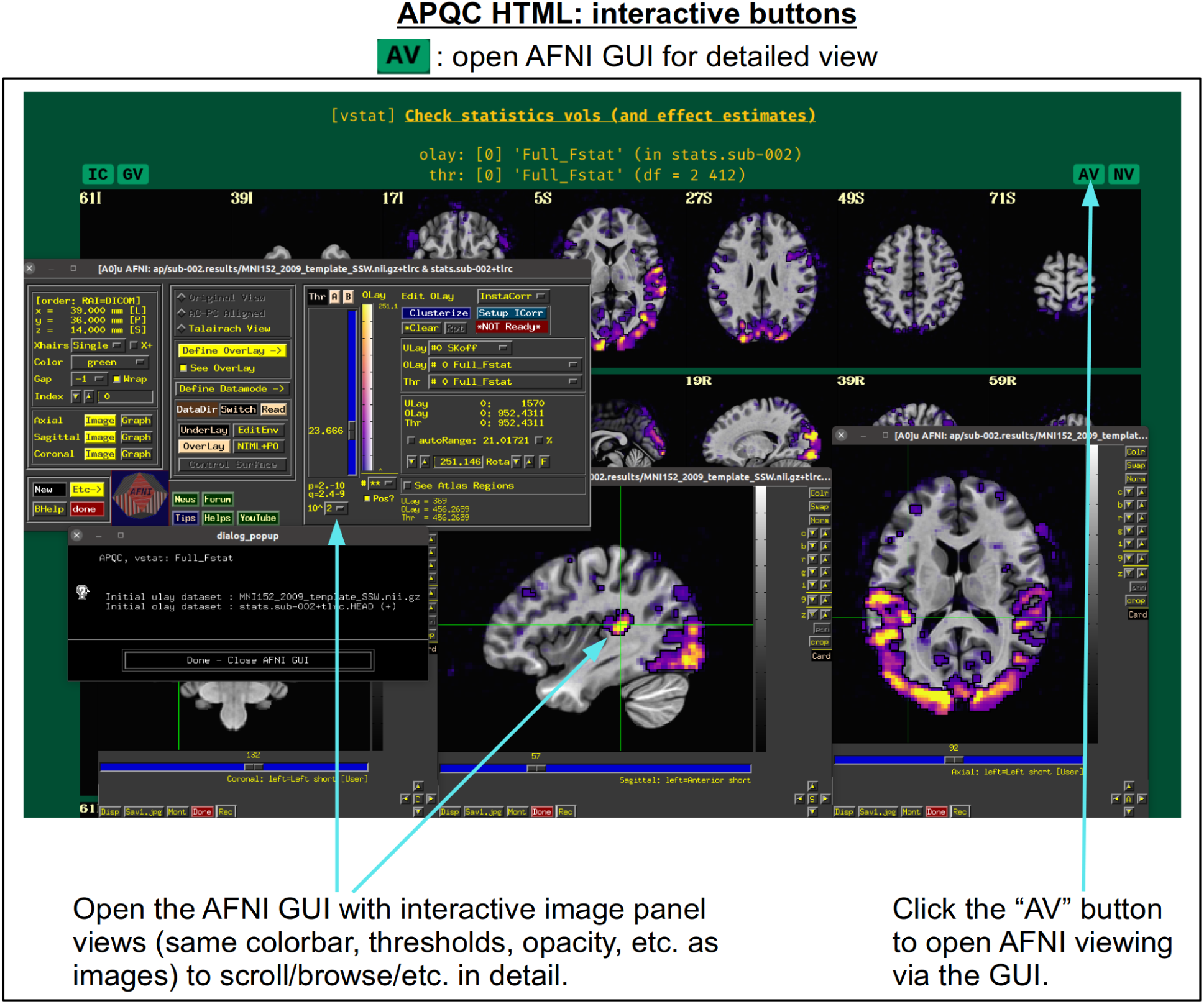
The "AV" button appears above several images within the APQC HTML. Clicking it will open the AFNI GUI with that image’s datasets loaded, also in montages and with the same colorbar, thresholding, etc. so that the user may interactively investigate the data beyond what the systematic (but static) images show.

**Figure 7.**
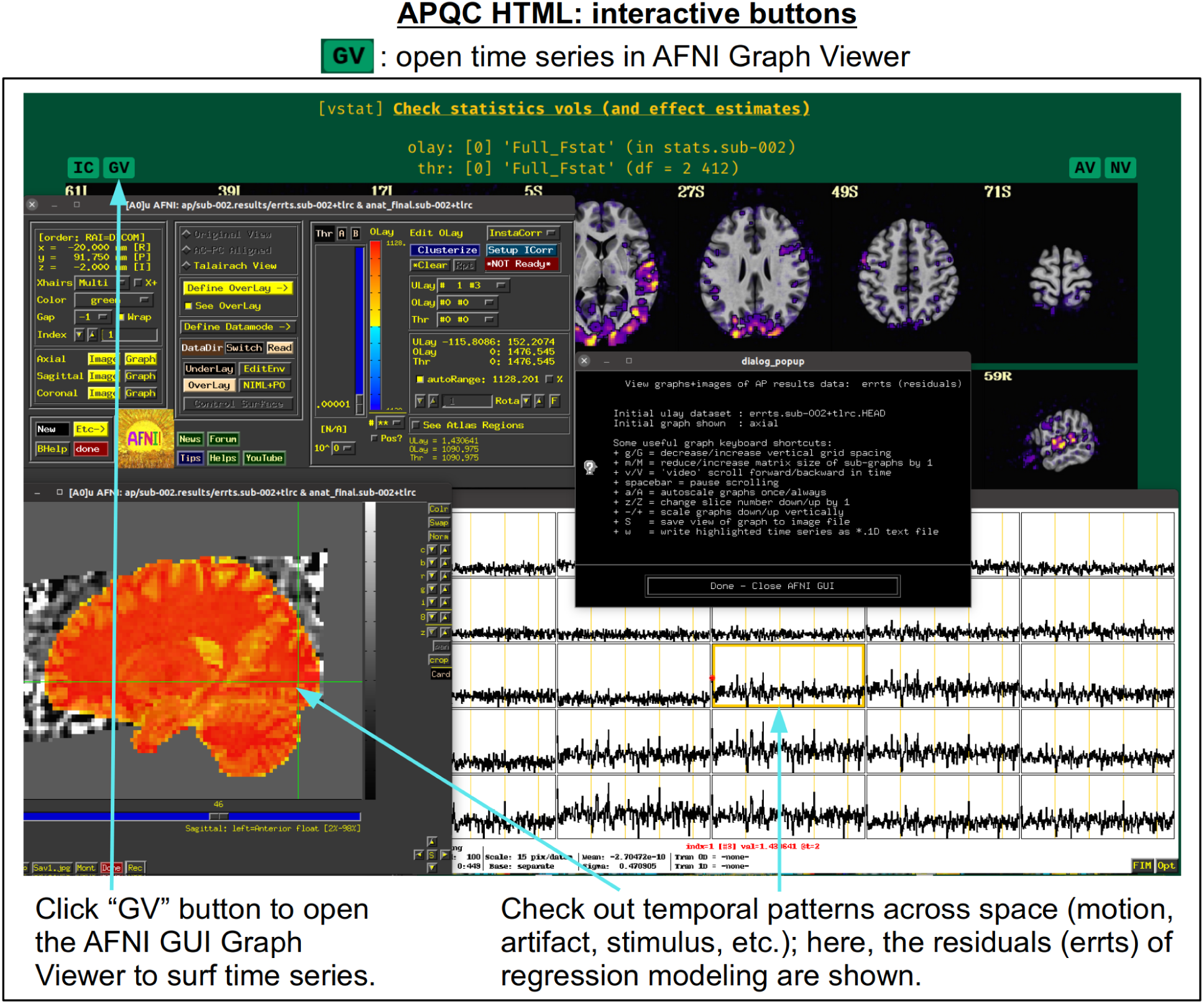
The "GV" button appears above several images within the APQC HTML. Clicking it will open the AFNI GUI’s Graph Viewer (as well as dataset image windows) with a time series dataset related to the data displayed in the image. This allows the user to interactively investigate the temporal properties around the brain, checking for odd features in individual time series or patterns across localized regions.

**Figure 8.**
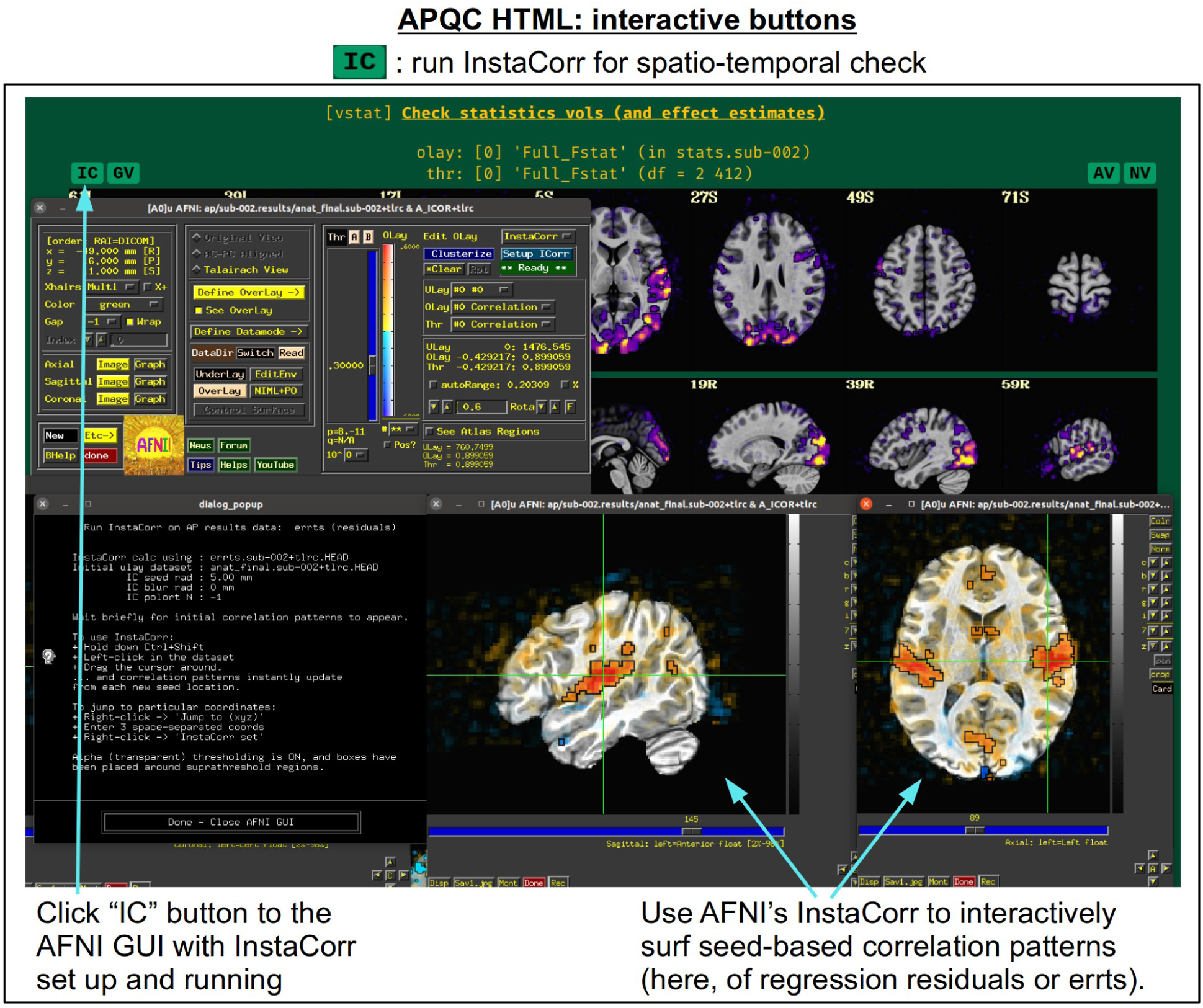
The "IC" button appears above several images within the APQC HTML. Clicking it will open the AFNI GUI with InstaCorr (Song et al., 2017) set up and running. With this functionality, a user can interactively investigate both local and global spatiotemporal properties: by holding down both the Ctrl and Shift keys, the user can click around the brain and see seed-based correlation maps that instantly update wherever they click. This GUI feature has been used many times to investigate EPI data for temporal artifacts. Having it now available and pre-loaded with a single button click from the APQC HTML greatly increases its efficiency.

**Figure 9.**
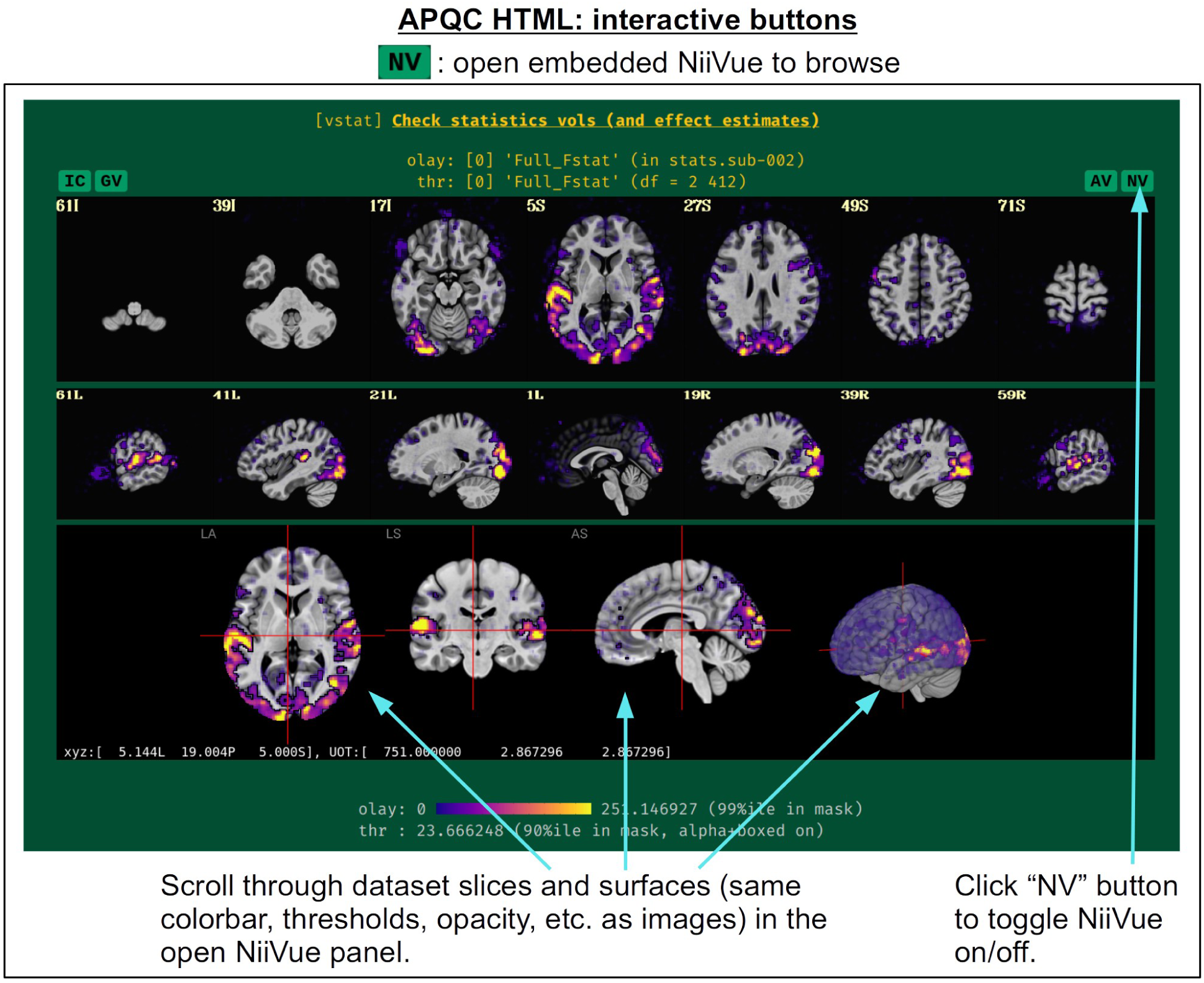
The "NV" button appears above several images within the APQC HTML. Clicking it will open a NiiVue instance (Hanayik et al., 2023) in a new panel within the HTML itself, loaded with the same underlay, overlay and/or threshold dataset and same coloration as shown in the image. Similar to clicking the "AV" button, the user can quickly and interactively surf through the data to examine any potential features or questions of interest. In addition to the three volumetric views, NiiVue also contains a 3D rendering, which can be rotated and even clipped (to see arbitrary interior slicings) using the mouse and keyboard.

These buttons enable such investigations directly from the APQC HTML.

For non-task datasets, such as resting state (Biswal et al., 1995) or naturalistic paradigms, a set of seed-based correlation maps are displayed if the template is in a recognized space. The default seed locations were chosen to be of interest in standard resting state (or naturalistic) paradigms, while also providing wide coverage of the brain with expected high correlation in general. The current seed locations are for the:

- default mode network (DMN), with the seed in the posterior cingulate cortex (PCC) and left hemisphere
- visual network, with the seed in the visual cortex and right hemisphere
- auditory network, with the seed in the auditory cortex and left hemisphere.

These seed locations have been defined in several template spaces, across age, geography and species. Currently, these include the MNI-2009c space (Fonov et al., 2011), the Talairach-Tournoux template (TT_N27), the various age ranges of the India Brain Templates (Holla et al., 2020), Haskins Pediatric Template (Molfese et al., 2021), NIMH Macaque Template v2 (Jung et al. 2021), and the Marmoset Brain Mapping v3 template (Liu et al., 2021). An example of the three network maps in the "vstat" QC block are shown in Fig. 10, for a resting state scan of a macaque that was aligned to the NMT v2 template as part of the MACAQUE_DEMO_REST demo dataset (see Jung et al., 2021). The known list of template spaces with preset seeds is expected to continue to grow.

**Figure 10.**
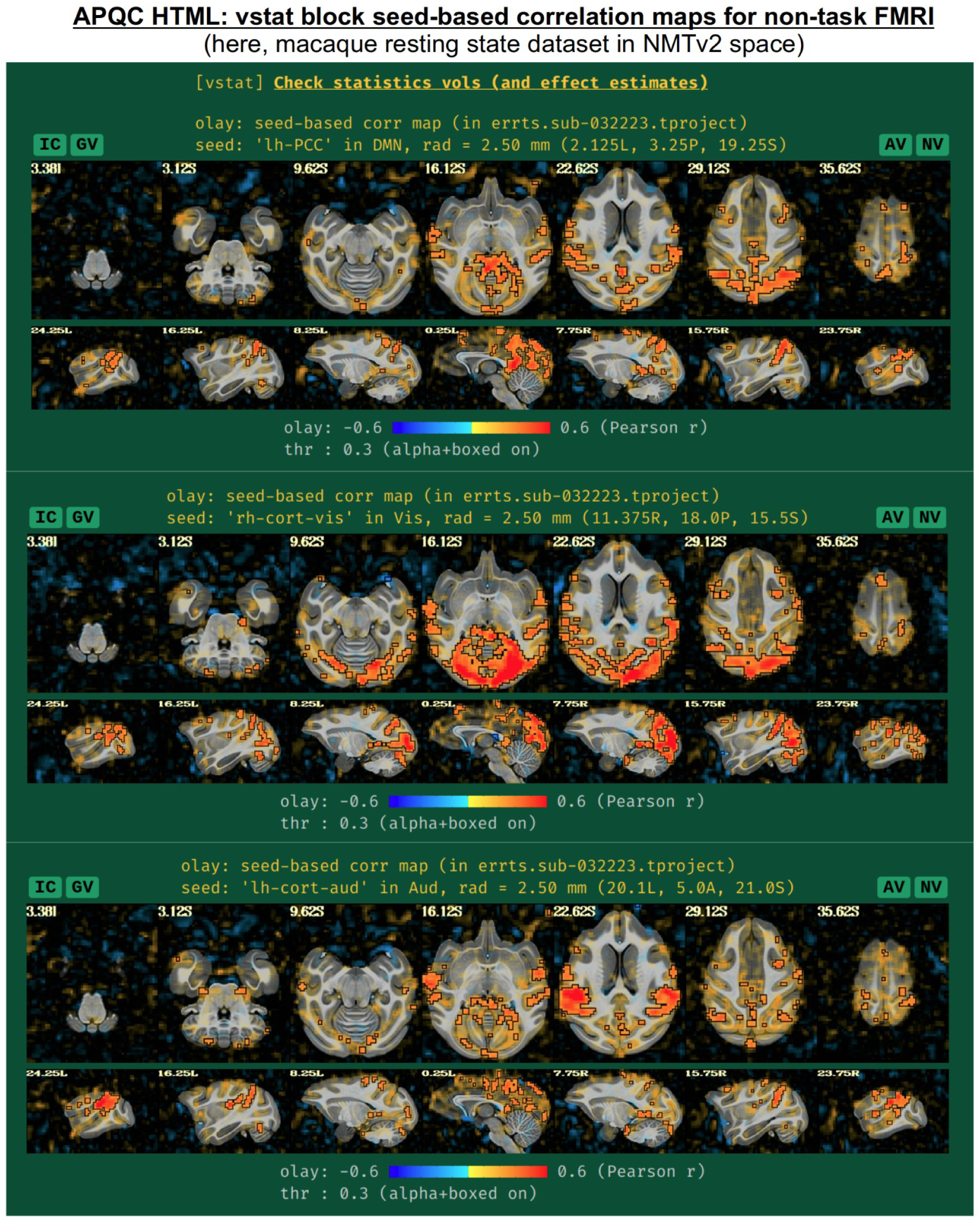
Example information within the "vstat" QC block, for the case of non-task-based FMRI. For known template spaces, three seed locations have been chosen to represent functional networks of general interest and wide brain coverage. These include the default mode, visual and auditory networks. These are shown here for a resting state macaque dataset that was aligned to the NMT v2 template; this data and full set of scripts can be downloaded by running "@Install_MACAQUE_DEMO_REST" (see details in Jung et al., 2021). The IC, GV, AV and NV buttons are part of the interactive APQC HTML functionality, in which the respective datasets can be opened in full GUIs if further examination is needed, described below in Figs. 6-9.

### mot: subject motion (including censoring and outlier detection)

Subject motion remains a major cause of concern in FMRI analysis and results reporting. The primary quantities for evaluating and thresholding motion events in AFNI are the "enorm" parameter (the Euclidean norm of the forward differences of the six rigid body motion parameters, estimated via volume registration^4^) and the "outlier fraction" (the fractional count of voxels in a brain mask that are outliers in their time series). These values are shown for all EPI datasets being processed, along with user-specified censor limits (Fig. 11); the duration of individual runs is shown by alternating the plot background between white and light gray. Red vertical bands show the location of any censored volumes (ones with suprathreshold enorm or outlier fraction values), and boxplot distributions of the parameters are also shown at the right, both before censoring (BC) and after censoring (AC). The fraction of censored volumes, as well as their index range selectors, are shown below the plots.

**Figure 11.**
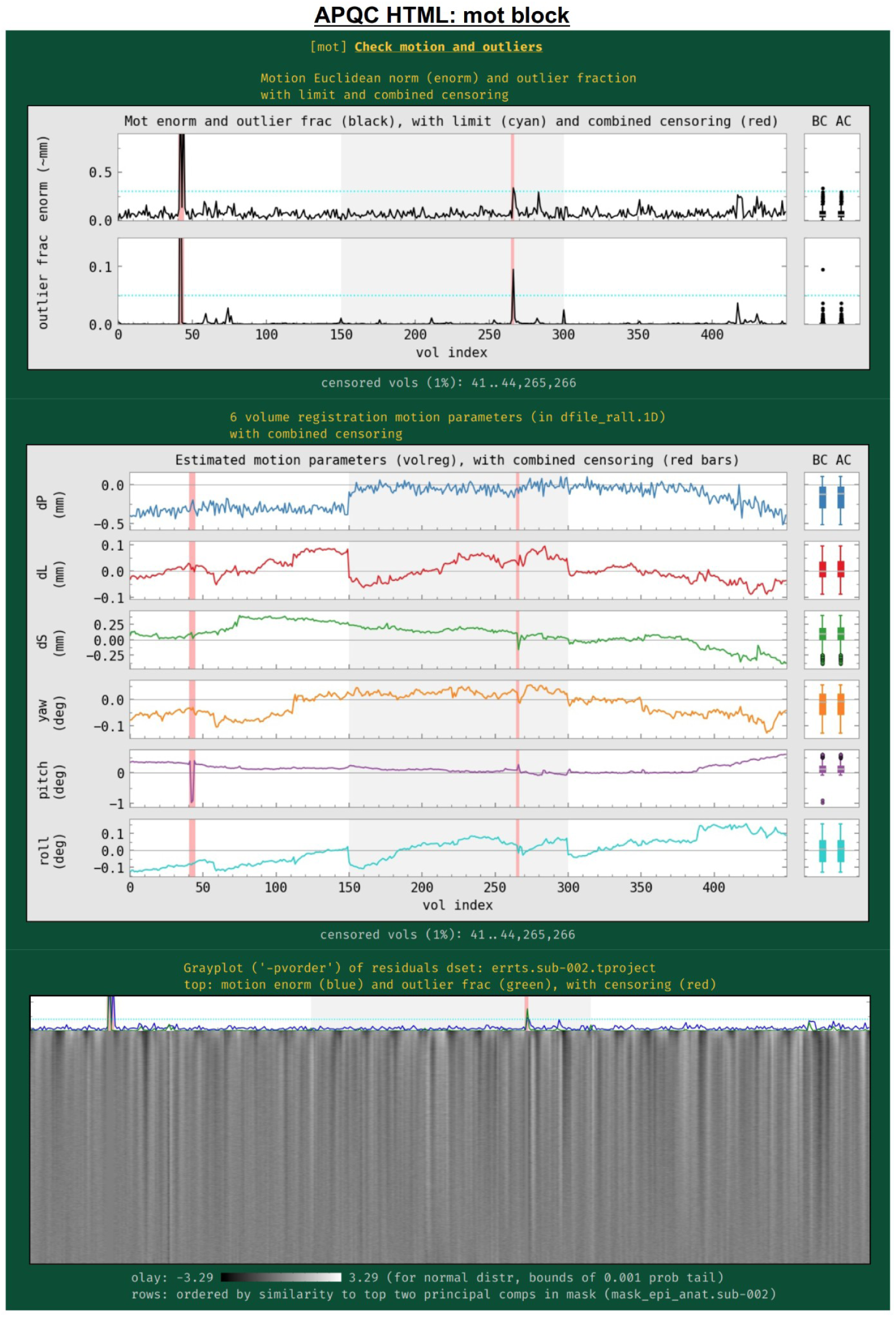
Example information within the "mot" QC block. Plots largely related to subject motion (as well as other to other features) are shown in this block. The between-volume motion parameter Enorm (Euclidean norm of 6 rigid body motion parameters) and within-mask outlier fractions are shown in the top row, along with censor thresholds (horizontal lines) and any censored time intervals (vertical red bands). BC and AC boxplots at the right show distributions of the values before and after censoring, respectively. The middle plot shows the individual parameter motion estimates, and the bottom panel shows a grayplot (where voxels have been ordered by similarity). Since afni_proc.py can process multiple EPI runs simultaneously, the background of the plots alternates between white and gray for each successive run, as shown here.

Also in this section (Fig. 11, middle), the time course history of the six estimated motion parameters themselves are shown. This can be useful as a follow-up to determine the underlying cause of motion— such as: yawning, nodding, pillow relaxation, ventilator, etc.—as well as of potentially apparent motion, which can be caused by respiration-induced magnetic field effects. These plots also show where censoring occurred, and have BC/AC boxplots.

Finally, a "gray plot" or "carpet plot" of FMRI time series across the brain can be useful for visualizing nonlocal/global features in time series, which can include motion (Power, 2017). In such plots (Fig. 11, bottom), each time series is then represented as a fluctuation in grayscale coloration, and a 2-dimensional matrix is created: time increases across columns, and each voxel occupies a different row. There are many ways to stack the time series, and here AFNI’s *3dGrayplot* orders them by similarity (and with decreasing similarity to the principal vector (PV) components in the WB mask). The resulting graphic highlights similar upswings and downswings in brightness (and hence of time series across space): the displayed values are Z-scores values of the time series, with the grayscale range chosen statistically to have min/max values of two-sided tails at *p*=0.001; in this way, the plots can be compared across subjects. Additionally, the subject motion parameter enorm and outlier fraction time series are plotted at the top, for reference.

### mecho: multi-echo FMRI combination

Typical FMRI datasets are acquired with a single echo per volume, but acquiring multi-echo FMRI (ME-FMRI) is becoming increasingly common, to boost local SNR and potentially reduce artifacts (see Kundu et al. (2017) and op cit). Within *afni_proc.py*, one can select many different forms of echo combination, including its own optimal combination (OC; Posse et al., 1999) calculation and the original MEICA algorithm (Kundu et al., 2012). More recently, it has integrated the MEICA calculation from the tedana software package (DuPre et al., 2021), and the APQC HTML directly includes that tool’s QC HTML report within its QC directory and as a link from the APQC HTML page. Fig. 12 shows an example of this, using data from AFNI’s APMULTI demo (which can be installed by running *@Install_APMULTI_Demo1_rest*; see Taylor et al., 2022).

**Figure 12.**
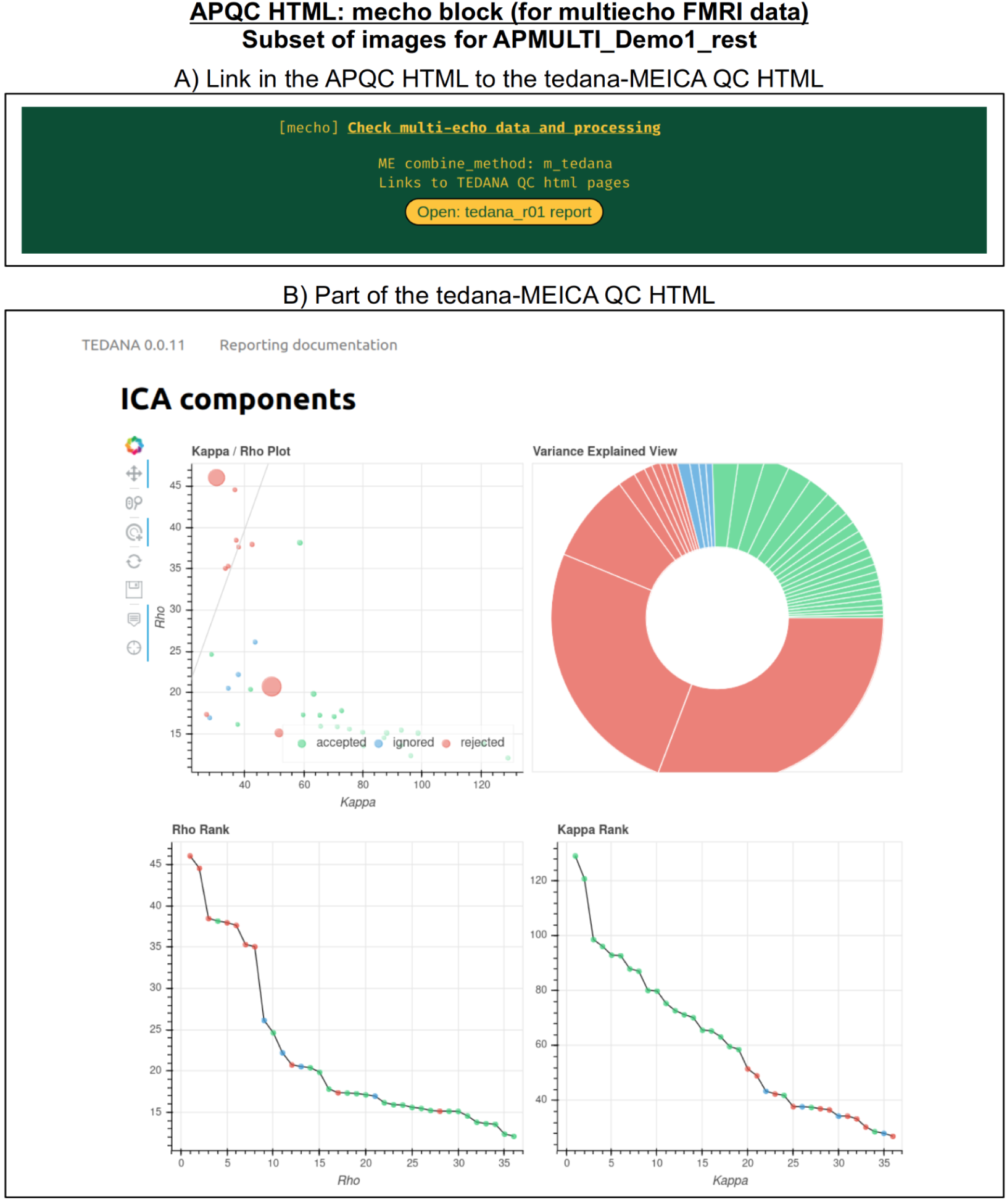
Example information within the "mecho" QC block, linking to the tedana software (DuPre et al., 2021) package’s QC HTML for MEICA, when that tool is used for echo combination within afni_proc.py. The top panel shows the link within the APQC HTML page, and the bottom panel shows the tedana QC output. A copy of the tedana HTML information is contained within afni_proc.py’s QC directory, so it remains portable within the APQC HTML.

### regr: regression and modeling information

In addition to showing the statistical results of modeling in the volume in the "vstat" block, the HTML also contains evaluations of additional regression modeling-related features, such as stimulus timing, spatiotemporal structure in the residuals, (or "error time series", errts), total degrees of freedom and TSNR. First, the regressors of interest are shown both individually and as their "summation" time series (Fig 13). These are the convolved time series, using whatever hemodynamic response function the researcher chose as appropriate for the task (blocks, events, amplitude modulation, etc.). This allows verification, for example, that the stimuli were well-spaced in time, and that a mistake such as entering the same file twice was not made in specifying the model (which is an all-too-common occurrence). Again, bands where censoring occurred are shown, providing a check on whether there might be task-correlated motion in the data (for each stimulus, the fraction of response time that has been censored is also calculated and automatically checked in the “warns” block, below).

**Figure 13.**
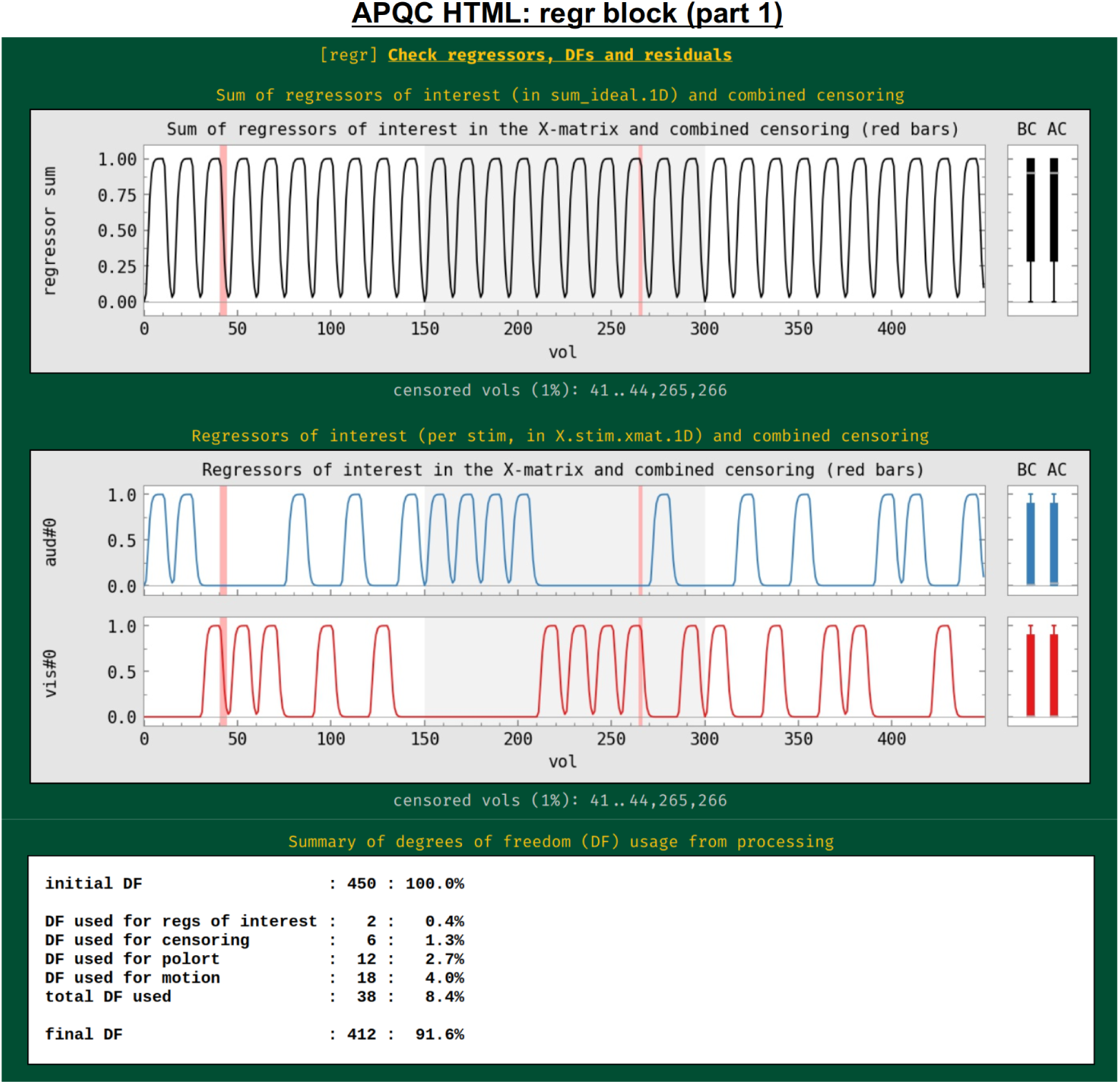
Example information within the "regr" QC block (and continued in Fig. 14). The top plots show the idealized stimulus responses (timing information convolved with appropriate hemodynamic or other response function) for this subject, first as the sum of all responses and then for each individual response. Intervals of censoring are similarly shown (see Fig. 11 caption for other line plot features). The table at the bottom shows the breakdown of degree of freedom (DF) usage within the subject’s regression model. Researchers must be careful to not use up too many degrees of freedom within their processing.

The process of model fitting is aimed at improving both the accuracy and precision of effect estimation, while also explaining as much of the variability in the input time series as possible.

The researcher adds regressors to the model with these aims, but these also reduce the degrees of freedom (DFs) in the associated statistics. Since researchers do not know a priori which regressors might only help the modeling, they must balance modeling considerations with the costs of including more regressors. For task FMRI, the translation of an effect’s *t*-statistic to a *p-*value significance depends directly on the remaining DFs. For non-task FMRI, the effective error bar of correlation values also depends directly on the remaining DFs. The breakdown of DF usage in the modeling are shown in this QC block: motion regressors, slow-drift regressors, censoring and bandpassing are common model components that reduce Dfs. This is an important and straightforward evaluation of any modeling process, even though it is often overlooked in many cases. For example, common "low frequency fluctuation" (LFF) bandpassing of resting state data to approx. 0.01-0.1 Hz is extremely expensive in terms of DFs: for EPI with TR=2s, this uses up almost 60% of the data’s DFs, and for TR=1s, this uses up nearly 80% of DFs.^5^ Because of this heavy reduction of DFs, a dataset may quickly become unreliable to use, depending on the amount of other regressors included and particularly the amount of censoring of subject motion (each time point of which also lessens the DF count by 1). Because of this fundamental issue, warnings about DF counts are included in the "warns" QC block, displayed below.

To further evaluate the modeling across the brain, the spatial correlation structure remaining in the residuals is shown in the corr_brain map (Fig. 14), which is the correlation of each voxel’s residual time series with the average time series of all residuals across the brain (which might be called the "global residual"). This image is transparently thresholded at |r|=0.3, with colorbar boundaries at |r|=0.6. Seeing large negative correlations or high-magnitude patterns that extend across non-GM tissues can often be signs here of problems, such as scanner issues, motion-related artifacts, or having too few DFs left in the time series.

**Figure 14.**
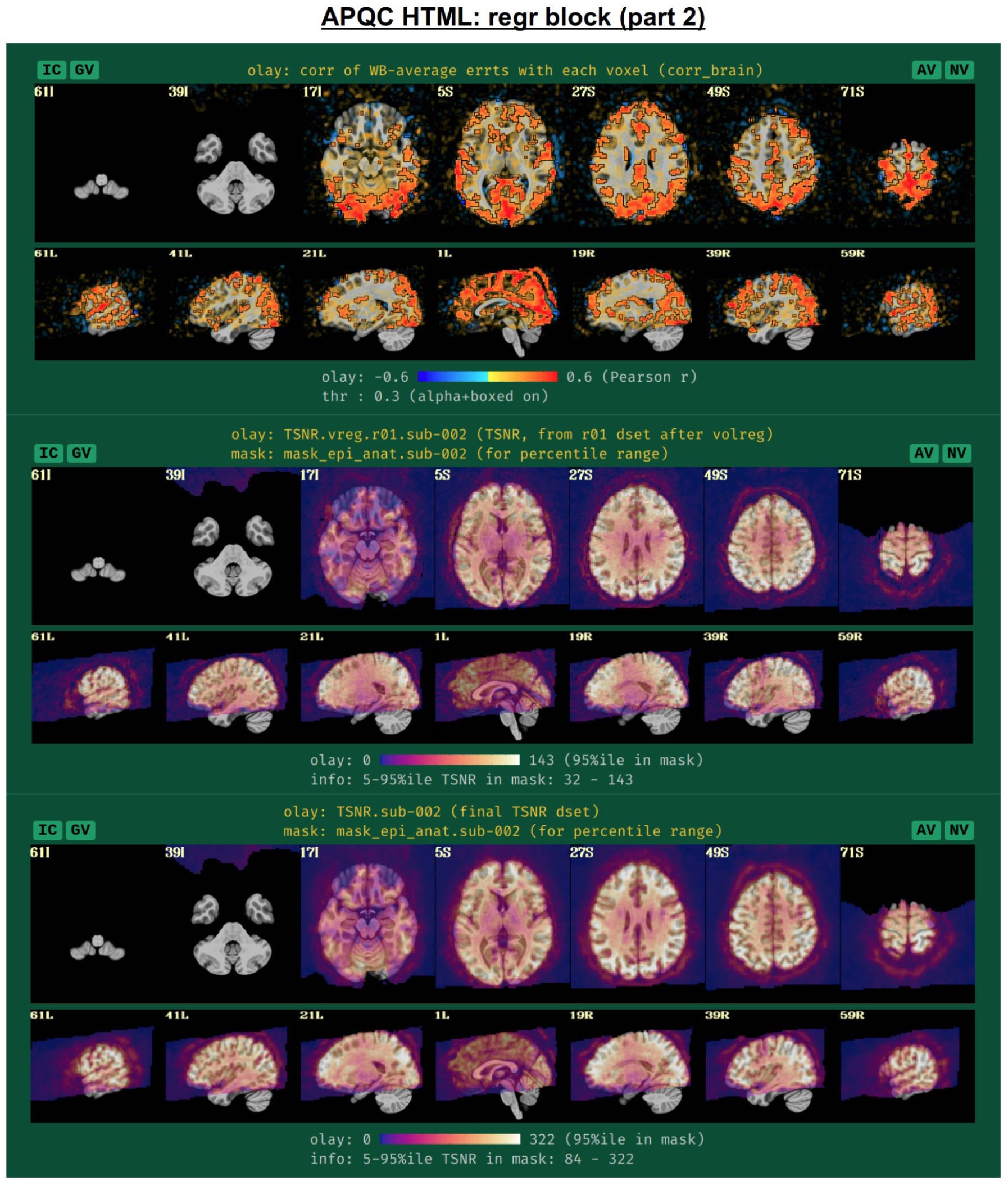
Example information within the "regr" QC block (continuing from Fig. 13, and see Fig. 15). The corr_brain map at the top shows the correlation of each voxel’s residual series with the whole brain average residual time series. High correlation outside of GM can point to various artifacts, for example. The TSNR plots in the bottom panels are calculated at different stages of processing (see definitions in the main text). Patterns of inhomogeneity are particularly useful for evaluating the data, such as noting that some parts of the brain (like VMPFC and subcortex) might have too low of signal strength to usefully interpret, even though they are within the FOV.

Another set of useful volumetric maps are the TSNR plots, which can be calculated at different points in time. While TSNR is generically "average(signal time series)/stdev(noise time series)", the definitions of what are the signal and noise time series vary at different stages of processing---and, perhaps surprisingly, different groups/software use different definitions of these even at equivalent points. In Fig. 14, TSNR is first calculated directly after volume registration (middle plot), and the "signal" is the (full) time series and the "noise" is the detrended time series; the second TSNR calculation comes after regression modeling (bottom plot), and the "signal" is the full (all_runs) time series and the "noise" is the errts time series. The colorbar ranges in each case typically reflect percentile values within a whole brain mask, and therefore can vary across subjects. While a popular quantity, it is difficult to define absolute ranges of interpretation for TSNR. The patterns of relative magnitude of TSNR can often be the most enlightening property of this quantity. For example in Fig. 14, the EPI data has inhomogeneous TSNR coverage, and regions of the ventromedial prefrontal cortex (VMPFC) and subcortical nuclei in particular have very low TSNR, perhaps rendering these regions outside the realm of valid analysis. Such an issue is a quite common one to observe in FMRI, due to sinus-related artifacts; viewing plots like these can help researchers avoid overinterpreting noise-dominated signals.

Often, the success of a study relies on having adequate signal strength in a particular set of regions of interest. Mean TSNR might be relatively high, but if there is poor signal within focal regions of interest (e.g., in the occipital lobe for a visual study, or in the amygdala and nucleus accumbens for a subcortical study), then one cannot reliably interpret results involving them. Similarly, one should be using data whose acquired voxel size provides a reasonable number of voxels within the region, for stability of estimation. Therefore, we have added a new data table to the APQC HTML with a table of TSNR and shape properties for regions around the brain (along with a central coordinate and label name, where known). A default set of regions from around the brain have been picked for several known spaces (see Fig. 15 for the APQC atlas for the MNI template used here), selecting regions of various sizes throughout the brain from popular templates. Potential issues within the ROI TSNR and shape table are automatically highlighted with warning levels. Currently checked quantitative properties include:

- Nvox, total number of voxels in ROI (warnings if quite small)
- Nzer, number zero-valued voxels (warnings as the fraction Nzer/Nvox increases)
- Dvox, the maximum depth of this ROI, counted in voxels (warnings as this decreases)
- Tmin, T25%, Tmed, T75%, Tmax, which are the min, max and quartile values of TSNR (warnings as T75% decreases, and as the slope of values (T75% -T25%)/Tmed increases)

**Figure 15.**
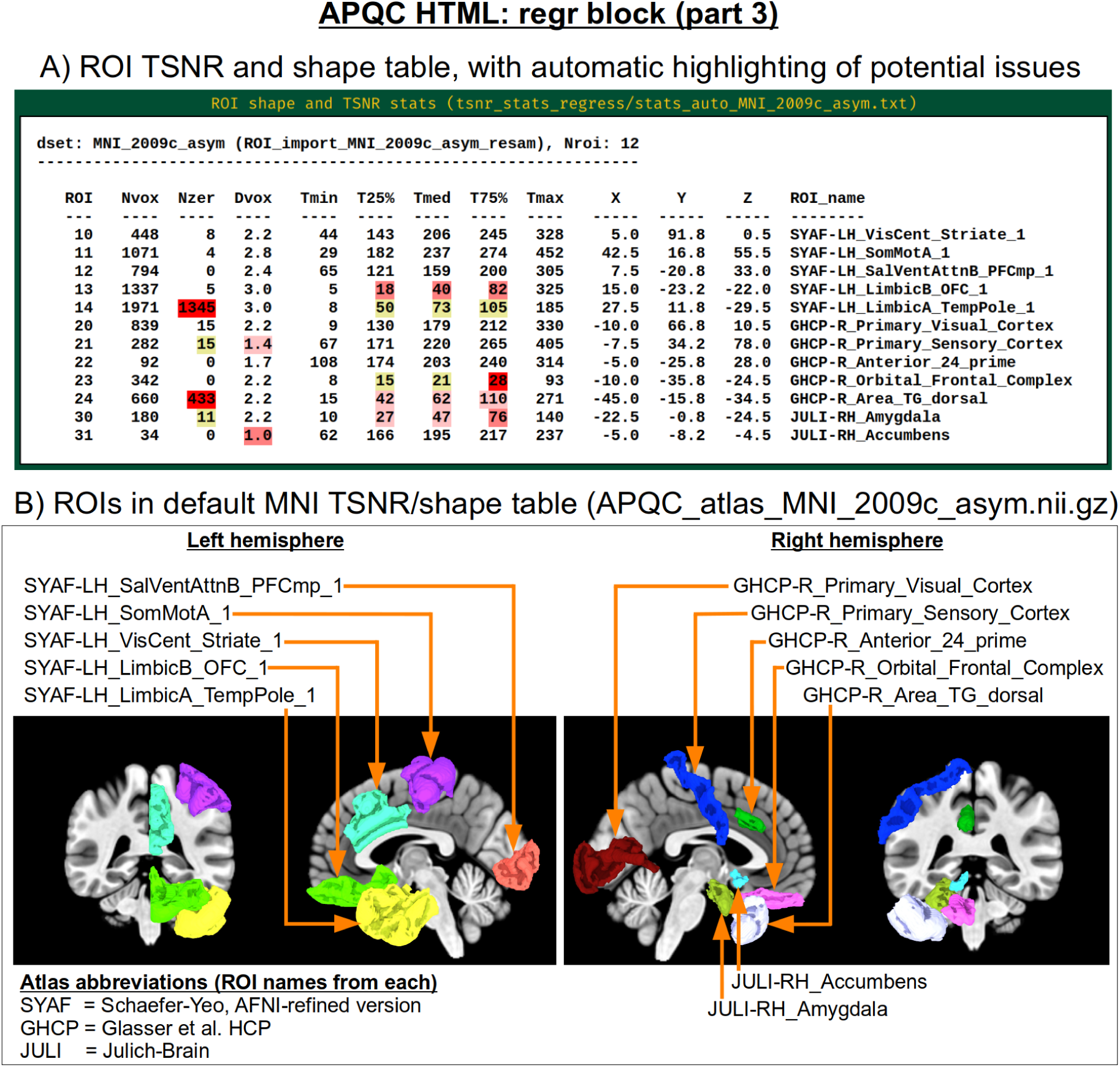
Example information within the "regr" QC block (continuing from Fig. 14). Panel A shows the ROI TSNR and shape table, with default ROIs for known template spaces (here, MNI-2009c, asymmetric). The table displays the following per-ROI information: number of voxels (Nvox), number of zero-valued voxels (Nzer), depth counted in voxels (Dvox), and then the minimum, 25%ile, median, 75%ile and maximum TSNR values. The coordinate of the maximum-depth location and the ROI name (stating first the atlas abbreviation) are also provided. Potential issues are highlighted with automatic coloring, such as too many zero-valued voxels or severe drop-off in TSNR, etc. as described in the main text. Panel B shows a SUMA rendering of the present MNI space ROIs. Where possible, different atlases are used per hemisphere to highlight choices based on either finer or coarser regions.

Researchers can also provide their own list of ROIs to be similarly checked for warnings, tailoring the QC evaluation to their particular study. Integrating these checks with pilot studies would inform researchers whether they need to change the way they acquire data to have more signal in areas of interest; it will also help those using public data to more efficiently know whether the data collection at hand is suitable for their research questions.

### radcor: using local correlation to check for scanner or motion-related artifacts

AFNI’s program *@radial_correlate* calculates the radial correlation (radcor) of each voxel in a dataset; that is, the correlation with a Gaussian weighted average of the voxels in a large radius (full width at half maximum (FWHM) of 20 mm, by default) around it. This was developed to detect certain kinds of coil artifacts, which can produce large, highly correlated patches across the brain. This map has also been found useful to detect patterns of motion that lead to large regions being highly correlated. These often appear as large patches of high correlation following the edge of the brain or EPI FOV.

In many cases checking the radcor values both before and after volume registration is instructive about the reduction of effects of subject motion in the time series (Fig. 16). Since *afni_proc.py* can process multiple EPI runs simultaneously, radcor is calculated for each run separately (except after the "regress" block, because then the multiple runs have been concatenated); this is reflected in Fig. 16, and only the images for the first runs are displayed for time series after *afni_proc.py*’s "tcat" and "volreg" processing blocks. Seeing high radcor values constrained within GM can be a positive indication that both motion effects are reduced (because those would likely produce correlations across tissue boundaries) and that task response is high (because that would likely increase correlations across functionally related GM). Note how the radcor patterns change, becoming less noisy and more constrained to GM, as processing occurs.

**Figure 16.**
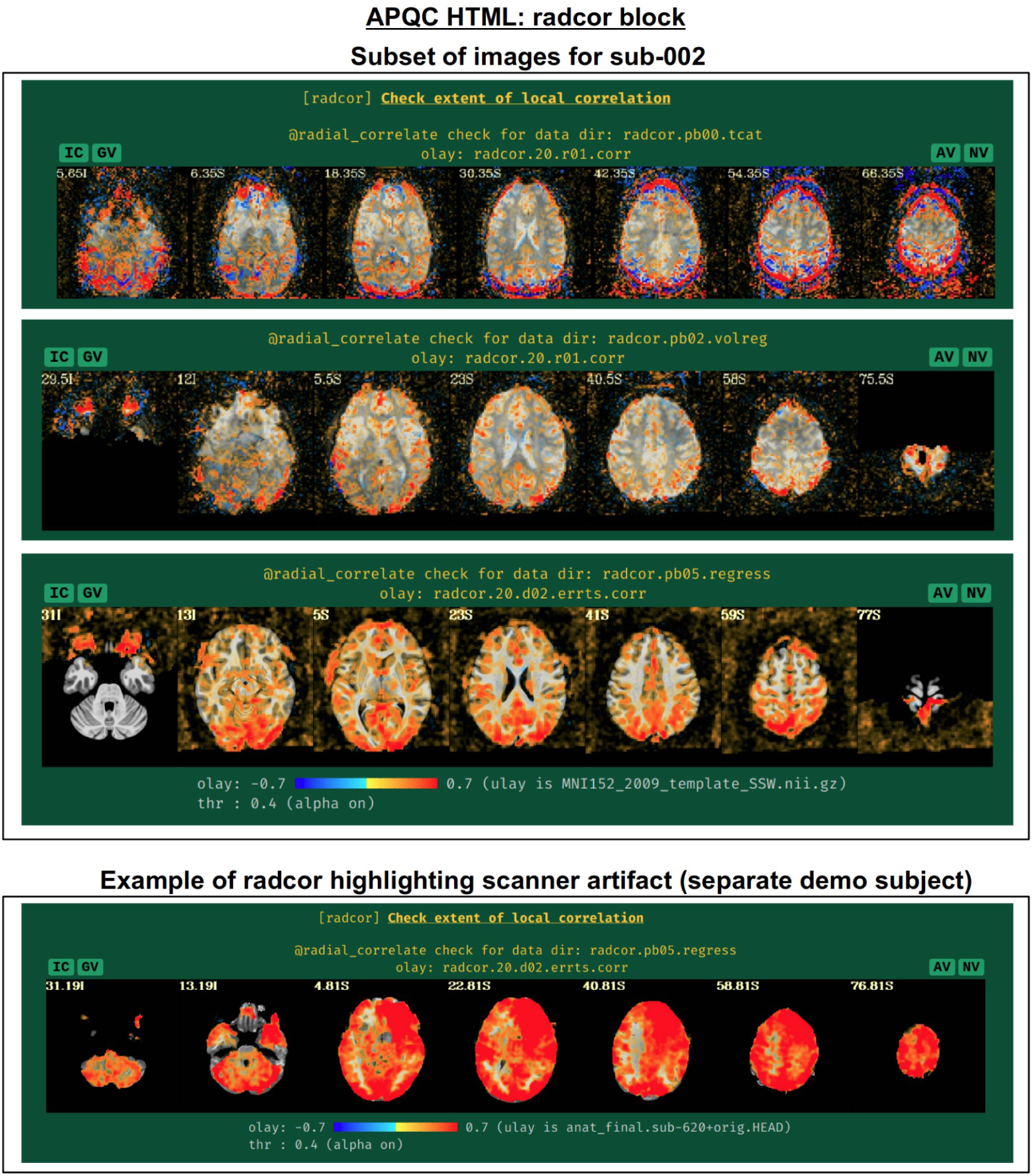
Example of the "radcor" QC block. The "radial correlation" (radcor) of time series highlights regions of the brain of high similarity. Reasonably high radcor might be expected in many GM regions, while high radcor values in WM and CSF or extending across tissue types might be signs of artifact. The radcor values typically change across processing: just after afni_proc.py’s "tcat" block (top panel), signatures of motion (ringing patterns of high positive and negative radcor around the edge of the brain) are visible; after the "volreg" block (second panel), where motion correction has been applied, these patterns are greatly reduced, and then further reduced after regression modeling (third panel). An example of an artifactual pattern, here highlighting a bad scanner coil, is visible for a different subject in the AFNI Bootcamp demo data (bottom row).

An example of a prominent scanner artifact visible in radcor plots is also shown in Fig. 16 (bottom panel). The anterior and right octant of the brain is extremely highly correlated; evaluating radial correlation here helped to identify a bad coil in the scanner. The researcher might view this data as unusable, or try applying a processing step like ANATICOR to reduce the artifactual features (Jo et al., 2010). This subject’s data comes from the AFNI Bootcamp demo download (AFNI_data5, s620_rest_c3r1+orig.HEAD).

### warns: internal checks

It was noted above that the *afni_proc.py*-generated script contains several internal checks on the data, outputting each result along the way. These potential warnings (see the subsection: *afni_proc.py output, II)* are collected here, along with images to help validate and provide more detailed information. For each warning, there is an assigned level with a color, and the maximum warning level’s color is displayed in the "warns" block QC label in the HTML’s top panel (Fig. 17): none (green), undecided (yellow), mild (light pink), medium (darker pink) and severe (bright red). The lowest three levels color the text of the word "warns" (leaving the background black), while the latter two invert the coloration for emphasis (text remains black, background colored). Importantly, the assigned levels are not absolute, but instead meant to be approximate guides and draw increasing attention to potential problems.

**Figure 17.**
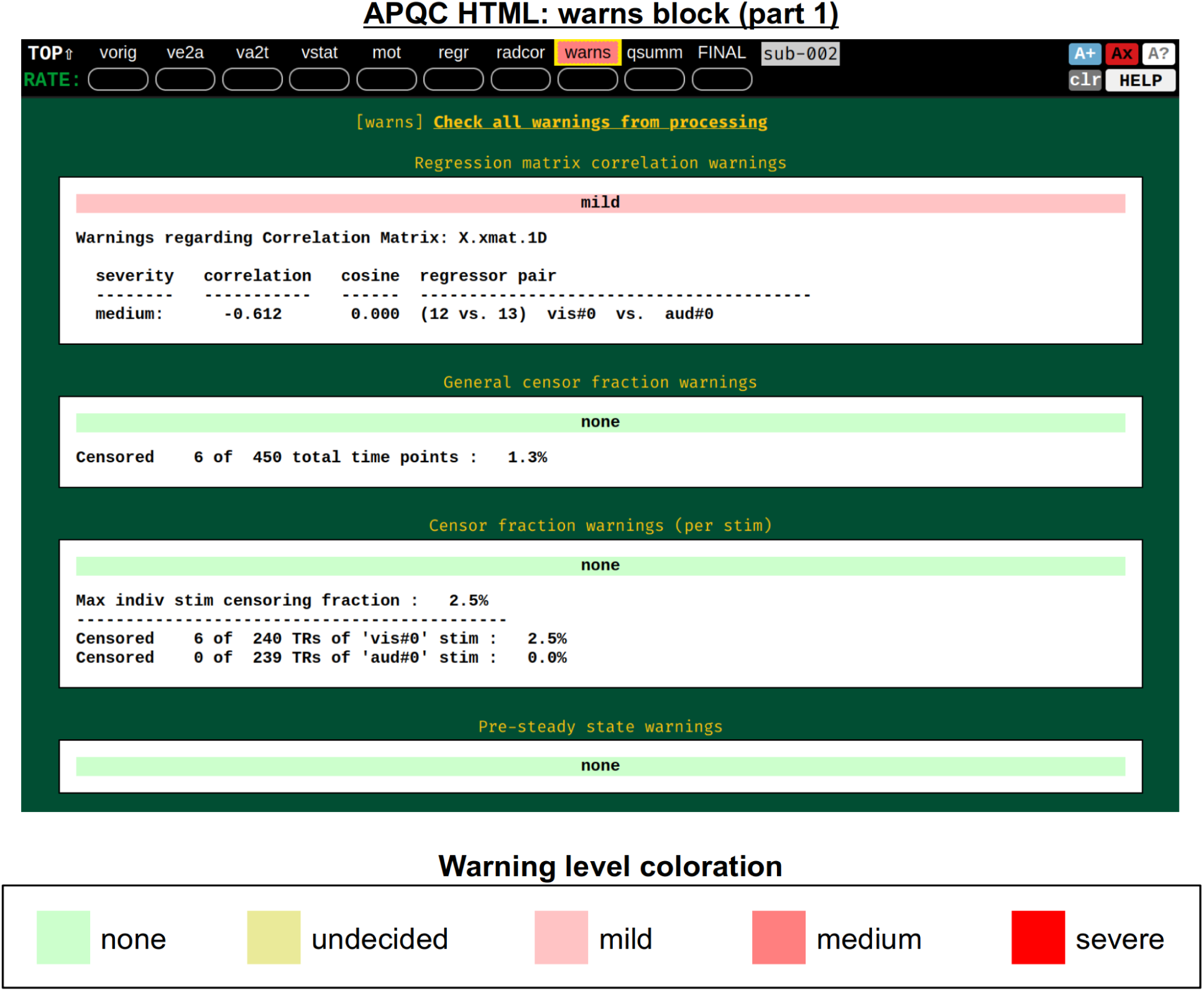
Example information within the "warns" QC block (and continued in Fig. 18). The automated text warnings from afni_proc.py’s processing script are accumulated in this QC block, with approximate warning levels assigned to each. The coloration of the most severe warning is applied to the "warns" label in the top menu; here, there is a "medium" level warning lower down within the QC block, shown in Fig. 18.

Fig. 18 shows two parts of the "warns" block that benefit greatly from having associated images to evaluate, in addition to the automated quantifications. The researcher can also add a flag to the *afni_proc.py* command to check for left-right flip mismatches between the EPI and anatomical datasets ("-align_opts_aea -check_flip"), following Glen et al. (2020). This is done quantitatively by comparing the cost function values for aligning the data "as is" with that of a left-right flipped EPI, and qualitatively by displaying the results (which are shown in the QC). This has been shown to be useful in several cases, even in widely used public repositories such as ABIDE, OpenNeuro and the Functional Connectome Project. The images help validate whether the estimated cost function comparison matches with the data (i.e., the cortical patterns of the chosen "correct" answer really do match better than in the other case); in fact, using AFNI’s typical local Pearson correlation (LPC; Saad et al., 2009) for this process, left-right flip guesses for human data tend to be highly accurate and were even useful in identifying a dataset whose EPI and anatomical appeared to erroneously belong to different subjects (Reynolds et al, 2023; Teves et al., 2023).

**Figure 18.**
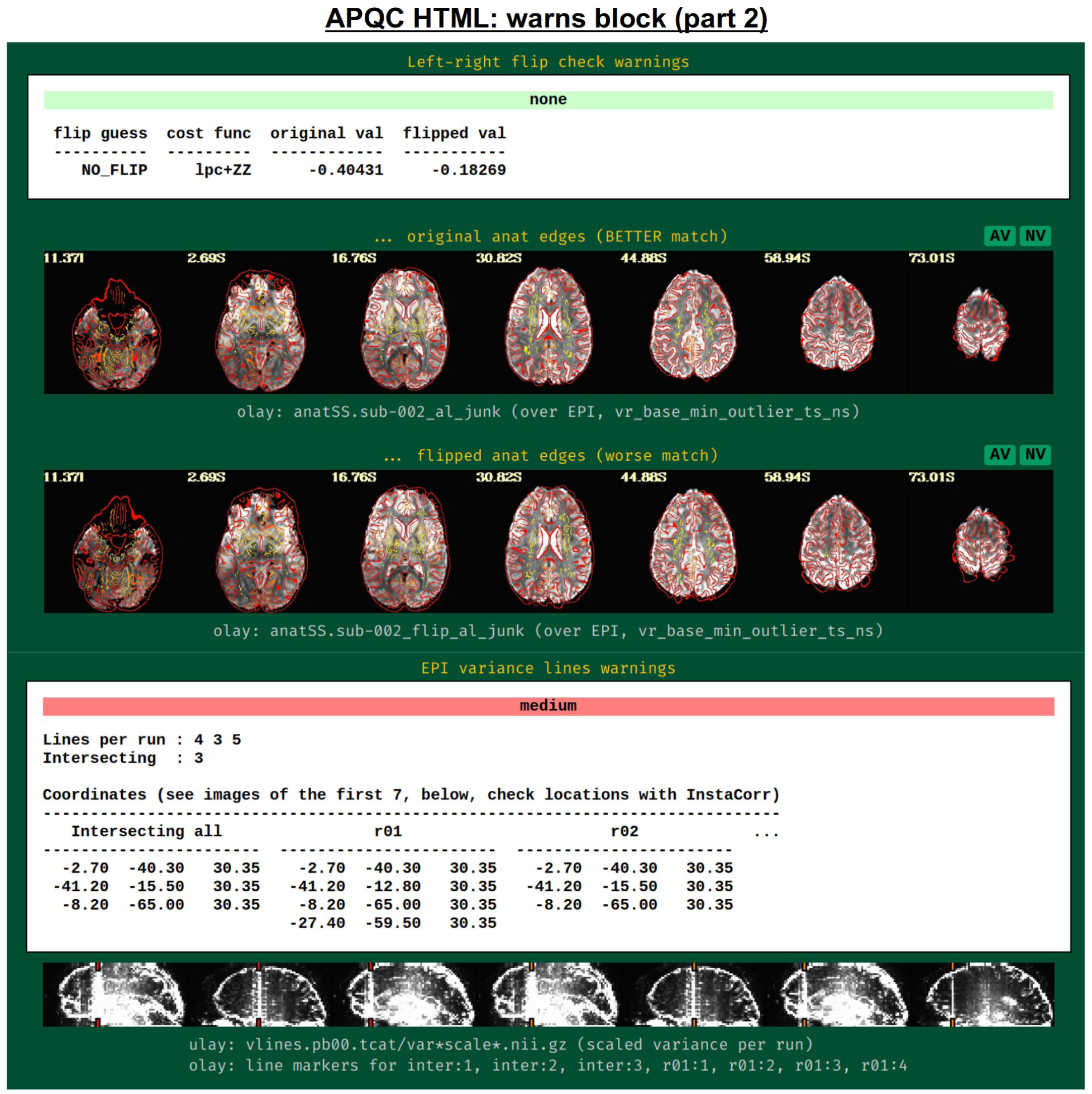
Example information within the "warns" QC block (continuing from Fig. 17). Both of the automatically evaluated warning items here benefit greatly from having associated images displayed. First, checking for left-right flips between the subject EPI and anatomical can be verified by looking at the cortical folding patterns; in humans, this check has tended to be highly accurate for reasonable quality data (Glen et al., 2020). In the lower section, EPI lines of high variance that occur vertically and narrowly have been observed in many data collections in public repositories; these can greatly affect correlation patterns and group analyses.

The bottom panel of Fig. 17 shows an automated check that was recently added to the APQC HTML. It was observed that many datasets, also in major public repositories, can contain narrow lines of high variance that are orthogonal to slice planes. In the FMRI Open QC project, this was observed through different means (see Reynolds et al., 2023), namely through noticing some odd TSNR patterns and then following up with InstaCorr in the AFNI GUI (that was also before the addition of the "IC" button within the APQC HTML). This automated check was specifically developed to more easily and automatically identify datasets that may contain this artifactual pattern. Because in practice complicated variance patterns can appear in EPI datasets, it is useful to have the images here to quickly check the patterns, as well as the IC, GV, and AV buttons to be able to follow-up interactively and efficiently.

Fig. 19 shows an example of this convenient functionality. The top row shows sagittal and coronal images after hitting the IC button (from the radcor block’s "tcat" image) and jumping to the first two EPI variance line table locations in the subject native space. The non-physiological correlation and anticorrelation patterns are obvious and notable in each case. Bottom two rows show that even after regression modeling, the signature patterns are visible (IC and GV buttons from the radcor block’s "regress" image): first using InstaCorr to find the equivalent locations in standard space, and then viewing the matrix of time series around that neighborhood where it is apparently that the baseline patterns and variability have different signatures in those corresponding vertical columns.

**Figure 19.**
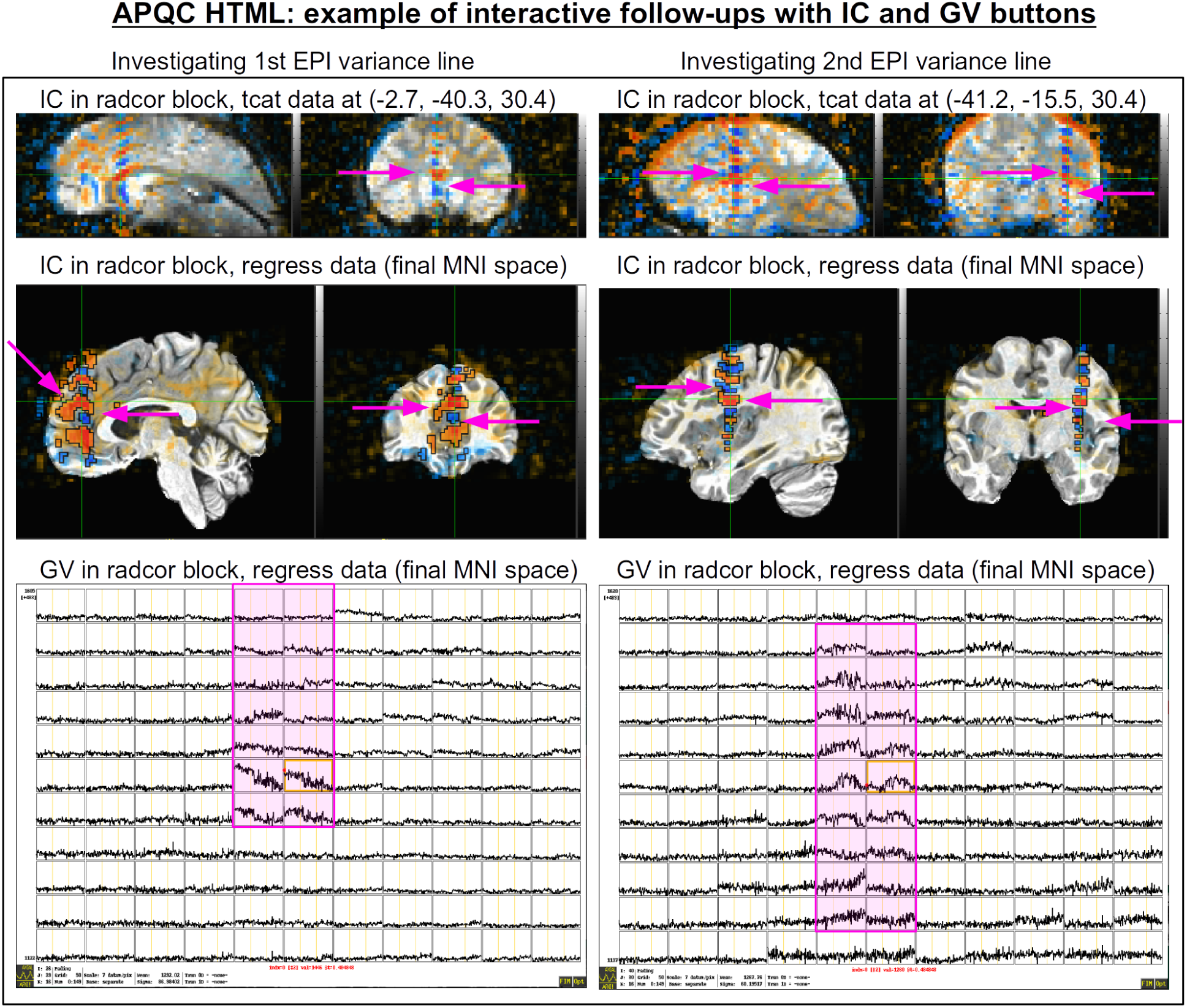
Example of efficiently and helpfully following up on variance line features using the APQC HTML’s embedded InstaCorr and Graph View buttons. (top) After clicking IC in the radcor block’s "tcat" image, one can jump to a coordinate location from the EPI variance line table and set the InstaCorr seed location in that subject’s native space. Patterns for the first two variance lines are shown in each of the respective columns. (middle) One can click on IC in the radcor block’s "regress" image and navigate to the equivalent anatomical location to view the artifact’s effect after regression modeling (and here, in standard space). It is still apparent, and (bottom row) clicking on the GV in that QC block shows the time series around that location: the vertical column corresponding to the strong correlation/anticorrelation pattern (highlighted in magenta) has obviously different patterns from the neighboring voxels.

### qsumm: quantitative summary of basic information

While processing, *afni_proc.py* keeps track of many basic quantities of interest relevant to the data and its processing. These include input dataset properties, processing-related metrics and derived summary quantities of interest such as mean TSNR and GCOR, as described above (see the "*afni_proc.py output, II*" subsection). This full set of quantities (i.e., what would be shown in the "ss_review_basic" output) are contained in this QC block. Researchers typically have a few "favorite" parameters to check for assessing their data sets; these can also be useful as exclusion/inclusion criteria when going to group level, as discussed in the next section.

### gen_ss_review_table.py: create group summary tables, subject drop lists and more

In addition to the subject-by-subject reviews described above, it can be useful to quickly summarize or tabulate quantitative results across a group. This can facilitate checking for patterns or trends that are helpful to understand when evaluating a data collection. Additionally, one might want to identify all subjects that have one or more of a particular set of properties, such as large motion or very low TSNR, for potentially determining which subjects to exclude from further study or other purposes. AFNI’s gen_ss_review_table.py program plays both of those roles, reading in one or more single subject (ss) dictionary files (either a JSON or colon-separated file, such as *afni_proc.py* creates) and both tabulating or filtering properties across that set of subjects.

At its simplest, gen_ss_review_table.py creates a comma separate variable (CSV) file, with each column being a quantity’s label and each row containing the respective values for one subject. A typical input to the command is the set of "out.ss_review.${subj}.txt" files, described above (and see Table 3) across all subjects. The created CSV file can be viewed as a standard spreadsheet and/or read into a program for further plotting, analysis or other comparisons. Users can also add any number of options to query properties among the table to report "outlier" subjects, including the following comparison operators:

- "VARY": a subject’s property varies from that of the first subject in the list; for example: *’AFNI version’ VARY*.
- "EQ": a subject property is equal to a particular value; for example: ’*flip guess’ EQ DO_FLIP*.
- "GT", "LT": the value of a property for a subject is greater than or less than, respectively, a particular value; for example: *’TSNR average’ LT 100*.
- "GE", "LE": the value of a property for a subject is "greater than or equal to" or "less than or equal to", respectively, a particular value; for example: *’average censored motion’ GE 0.1*.

When reporting outliers, the ID of each subject whose comparison evaluates to "True" for one or more queries is displayed, along with the values that evaluated as such. One can also add a "SHOW" keyword to additionally display some other property for each subject in the "outlier" table, for additional information. By adding the "-show_keepers" option, users can essentially invert the set of subjects shown in the displayed table: each subject with *no* comparisons that evaluate to True would be shown. Some example commands, using quantities stored in the "out.ss_review.${subj}.txt" text files, are provided in Table 5.

**Table 5.**
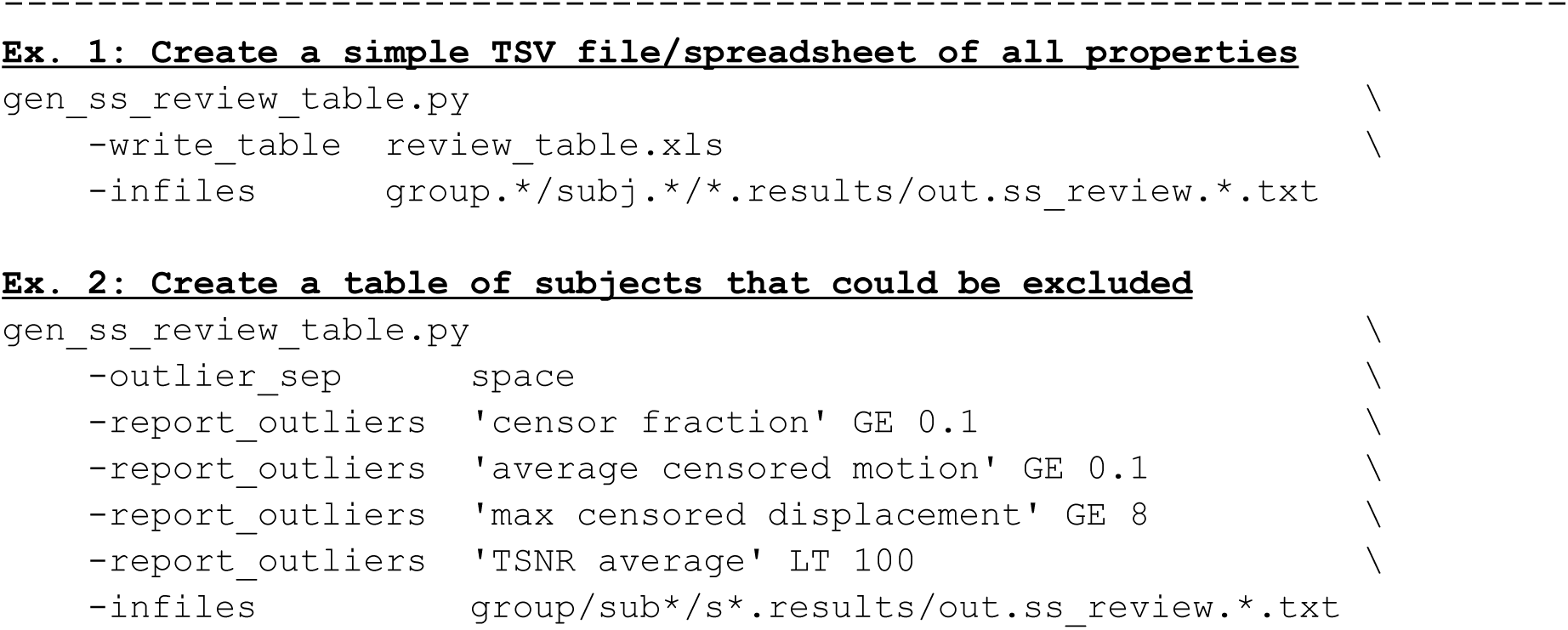
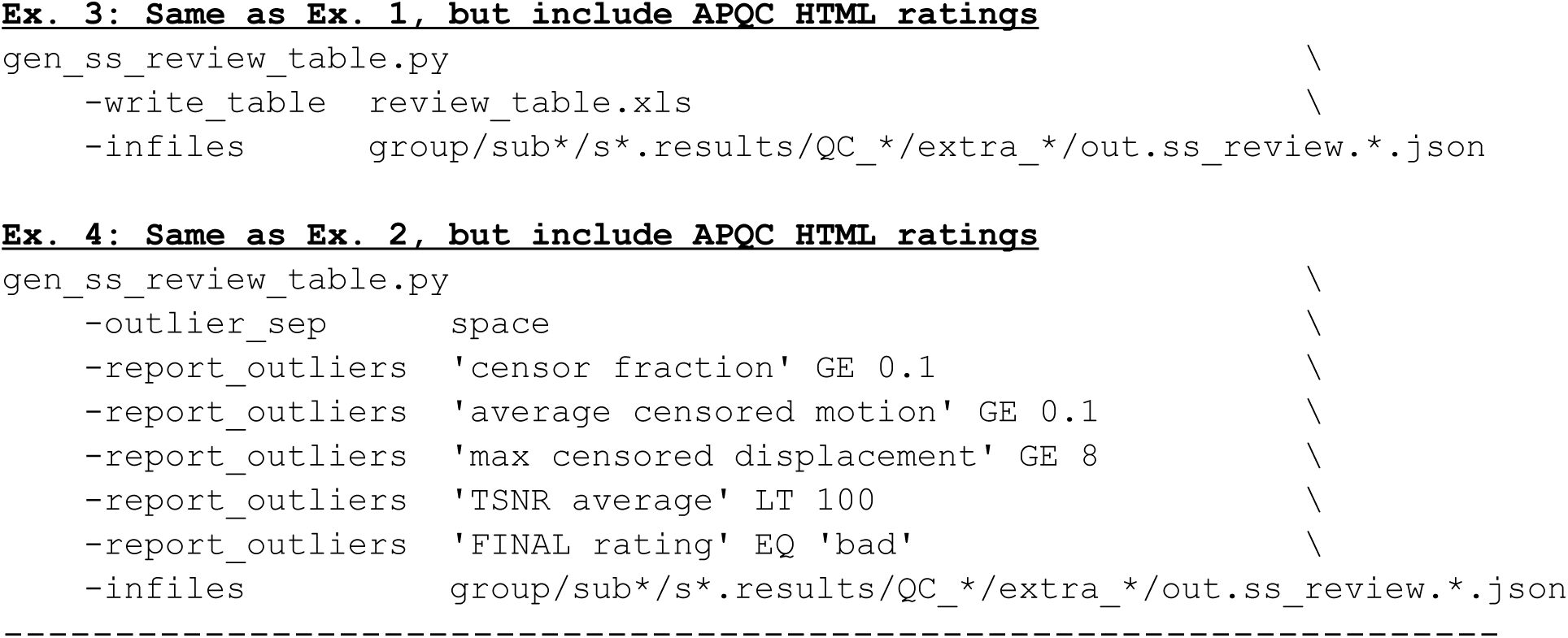
Examples of gen_ss_review_table.py commands that can be used to create tables of important quantitative and qualitative features across a group. By adding "-report_outlier" options to the command, one can apply one or more filtering criteria to display just the IDs of subjects with at least one chosen property (and the values of the properties that put the subject into the list are also displayed). By adding "-show_keepers", the table of subjects can be inverted. The set of input files are relatively flexible, including simply formatted colon-separated files and JSON files. One might commonly input the set of "out.ss_review.${subj}.txt" files created by afni_proc.py, or the set of "out.ss_review.${subj}.json" files in the QC directory; the latter are particularly useful after evaluating the APQC HTML, since the QC ratings are also added to the JSON file. Note that the APQC HTML rating values translate as follows (see Ex. 4 in the table): ’good’ (’+’), ’bad’ (’X’), ’other’ (’?’).

Beyond using the ss_review file calculated and stored by *afni_proc.py*, one can also use a version of that file that exists in the QC directory. When the QC_${subj} directory is first made, a copy of that information is put into a subdirectory, called extra_info/out.ss_review.${subj}.json; all the keys and values from the file in the results directory match exactly. The QC ratings that the user adds to the APQC HTML are also appended to this latter JSON file. Therefore, one can use gen_ss_review_table.py to directly combine the information from both the quantitative output file and the qualitative QC review into a single summary table, as well as apply exclusion criteria across a group based on that full set of quantities. This greatly facilitates the full reproducibility and shareability of drop criteria, within a single consistent structure.

### gtkyd_check: getting to know your data (and its consistency) from the beginning

Being able to compare lists or dictionaries of properties across files is also useful before processing even begins. For example, with EPI data one wants to know if matrix size, voxel dimensions and TR are consistent across a collection, as well as whether slice timing is included, etc. In Reynolds et al. (2023), checking these features was referred to as "getting to know your data" (GTKYD), and the authors noted a surprising degree of variability in fundamental properties within some public data collections.

To both simplify the running of this process and expand its output, we have since created a program called *gtkyd_check*, which first loops over a set of provided files (e.g., subject anatomical or EPI datasets) and creates a simple dictionary file of basic properties for each dataset. It then uses gen_ss_review_table.py to tabulate these into a spreadsheet (one dataset per row), and then again to create a "variability report" to quickly display any ones that vary. This provides a simple way to get to know one’s data quickly and to quantitatively evaluate its properties from the start.

In many cases, feature variability in raw data might represent a quality control issue about some datasets’ suitability for inclusion in further analysis from the start. Variability can occur due to scanner upgrades, console-typos, DICOM header conversion, incorrect mixing of datasets and more. When utilizing public data, one should always evaluate the fundamental data properties oneself, since the acquisition was performed by others and expectations/assumptions can vary across groups. We also view *gtkyd_check* as a valuable tool when acquiring one’s own data, again due to the practical realities and difficulties with the process. It can efficiently help to verify consistency (or to identify inconsistency) in a protocol as new datasets are acquired, which is the best time to check.

## DISCUSSION

FMRI data collection and analysis are each complicated. There are many things that can easily go wrong: coil issues, scanner console typos, file conversion errors, mistakes in selecting analysis parameters, and more. Additionally, subtle issues such as the omnipresence of magnetic field imperfections, unavoidable distortions (B0 inhomogeneity, sinus-related signal loss, etc.), and subject motion can create interwoven issues that in many cases belie simple interpretation as "good" or "bad", but that must still be understood and dealt with. The process of performing quality control on data should encompass more than simple box-ticking of "include" or "exclude" a subject, and instead lead to much more complete understanding of the data in all of its stages from collection through intermediate (pre)processing to final analyses (Reynolds et al., 2023). As such, it should be actively planned during the study design phase (Teves et al., 2023), and some measures can be gathered and assessed even during the acquisition time (Etzel, 2023). QC checks remain necessary for evaluating the suitability of the data for the research at hand, and they should be integrated early on in data collection—preferably in a piloting stage. Ignoring these steps risks unfortunate and costly data waste (as well as participant time, scanner resources, researcher effort and more).

Many tools have been developed within the AFNI toolbox to facilitate this process of evaluating and understanding data, and we have described several of them here. These include outputting automatic checks and quantitative summaries, as well as facilitating systematic qualitative review with scripts and *afni_proc.py*’s comprehensive APQC HTML report. The latter in particular provides one place for examining a wide spectrum of information about the subject’s data and the results of the many processing steps: raw data, alignment processes, statistical modeling, motion estimation (and censoring), and more. Having a single HTML page with all of this information makes it possible to jump between QC blocks and obtain a fuller picture of the data than seeing parts in isolation—this is quite useful when diagnosing FMRI, given the often tangled nature of underlying issues. For example, if the statistical results are not as expected (in the "vstat" block), is it because the subject moved a lot (checked by jumping to the "mot" block) or because of a coding mistake with specifying stimuli files (just to "regr" block) or perhaps because of stimulus-correlated motion (also in the "regr" block)? This understanding can feed back to decisions about improving acquisition, updating processing choices, or filtering subject datasets.

The APQC HTML creation depends only upon a small number of open source tools (AFNI and a single, widely used and relatively stable Python module, matplotlib), without dependencies on licensed software; visualizing the QC is likewise straightforward, requiring only a modern browser without any other add-ons. The interactive features of the HTML depend additionally on only Flask and Flask-Cors, with almost no version restriction, further facilitating its portability. The recently developed NiiVue viewer (Hanayik et al., 2023) is now included within AFNI distributions. It has been developed to integrate many of the features used in the AFNI GUI (separate overlay and threshold selection, transparent thresholding and boxing, colorbars, etc.), creating a seamless integration in the QC process. Being able to view the data interactively within the single HTML (and to toggle the view off when done) is particularly efficient for following-up on potential questions quickly; the volumetric rendering, which can be sliced and scrolled through arbitrarily with keypresses, further adds to this capability.

The specific content of the APQC HTMLs follows from the researcher’s processing choices in running afni_proc.py. As FMRI datasets and study paradigms vary widely, there are many sets of options for running this command. A link for the GitHub code repository for the present paper, which includes an afni_proc.py command, has been provided above. For a more detailed presentation of afni_proc.py commands and ranges of options, see Reynolds et al. (2024) and Taylor et al. (2018). Additionally, the AFNI Codex^6^ provides an open, online resource of code examples associated with papers and projects, many of which contain afni_proc.py examples. There are several available demos of both human and nonhuman FMRI processing that are freely available to down with @Install_* commands in AFNI.^7^ The publicly available AFNI Bootcamp teaching material contains several examples of afni_proc.py commands, as well as gen_ss_review_table.py examples. There are several online tutorials for processing, including the AFNI Academy Youtube channel and recordings of previous Bootcamps.^8^

In complement to the APQC HTML, the *gen_ss_review_table.py* command provides a scriptable and reproducible way to combine the information from single subject processing to group analysis. This functionality can be used for creating complete summary tables, checking how the data from new subjects compare with earlier ones, or even listing subjects to filter. It is easily reportable within publications, making subject exclusion criteria programmatically clear. Moreover, it is easy to rerun and to recalculate tables if, for example, new subjects are added to the study. While the primary focus of this tool is for navigating quantitative features of the data, we have described how it also integrates the qualitative understanding through the tabulating QC ratings.

### The roles of qualitative and quantitative QC

In quality control—or, more generally, in the procedure of understanding data—there is an important interplay of qualitative and quantitative information, which are respectively based on visualization and on measures. Both are useful, and in best case scenarios they reinforce each other. Primary understanding of FMRI data originates with visualization, as it provides a rich assessment at multiple scales. Measures and metrics can summarize individual aspects of the data in different ways; as simple values, they can be queried and compared relatively easily. Typically, the most useful quantities are those derived directly from visualization, as they can be meaningfully calibrated or directly associated with artifacts. Thus over time, having more visualization and deeper understanding of data can foster the development of more useful measures. For example, GCOR, the left-right flipping metric and EPI variance line estimates are all quantities in AFNI’s QC that have been derived from visualizing data. However, given the complicated and multiscale nature of neuroimaging datasets, qualitative assessments are still needed to fully interpret scenarios (e.g., is correlation low because of poor alignment, large noise, low local SNR, or some other factor?).

There are still several processing steps that essentially require a researcher to assess them visually. For example, consider the quality of data alignment: the cost function values *are* the computer-generated metric of similarity, so they cannot feasibly be used for independent self-assessment. By definition any alignment that completes without error has reached its "internal" measure of requisite alignment quality, but that algorithm may have gotten stuck in a local minimum or simply not be adequate in practice.

However, the quality of alignment can be assessed nearly instantaneously by the human eye. In other words, automated methods are driven by objective cost functions, and visual inspection is specifically required to know if these metrics have been trapped by a local minimum or other unsatisfactory solutions. While work will certainly continue to increase the objectivity and data-driven aspect of FMRI QC within the field, at present (and for the foreseeable future) the informed analyst still seems needed for this important aspect of processing. The goal of QC is ultimately not just to "find subjects with different metrics", but instead to be (more) certain that there are no surprises or artifacts in the data. The goal of AFNI’s APQC HTML is to provide an organized, systematic and comprehensive means of evaluating and recording the quality of data processing.

The argument is sometimes made that QC should depend solely on quantitative measures, to make the process more "data driven", automated (for efficiency), objective (unlike subjective interpretations of images) and "unbiased". This implicitly assumes that a set of complete measures exists to fully evaluate FMRI data and their processing for quality, to automatically distinguish between the degeneracy scenarios that could produce a given set of quantities. However, at present such a set does not exist for FMRI neuroimaging: the field has a long way to go to first understand the data itself more completely, to be able to develop a broader set of measures. Consider automatic comparison of seed-based correlation maps: a measure of similarity is easily calculated, but many different situations could lead to a given value, requiring a human to interpret what might be happening. Visualization is still very necessary to (better) interpret features of the data, and to be able to use the QC information to make decisions about the data, its properties and its processing.

Finally, related to its role with fostering an understanding of particular datasets, visualization greatly facilitates education and training and building a more general understanding of data. This is particularly the case when images are made systematically and can be compared easily across subjects (Lepping et al., 2023). Integrated qualitative-quantitative reviews also usefully inform discussions of inter-rater variability, which necessarily exist in data assessment (Williams et al., 2023): consensus building becomes easier during the training phase, and differences of interpretation are more readily discussed. The APQC HTML usefully contributes to all of these aspects.

### Integrating QC in studies

There are many points at which quality control can (and likely should) be integrated within a study. An example schematic diagram for applying the primary tools described here is shown in Fig. 20. Similar considerations apply whether one is acquiring one’s own data or using data collected elsewhere, such as when downloading from an online repository. The only real difference in those two situations is that in the first case, QC will be applied in a rolling manner as datasets come in—it should never be delayed until after a large amount of datasets have been acquired, since there is a danger that then a large set will have issues and need to be wasted.

**Figure 20.**
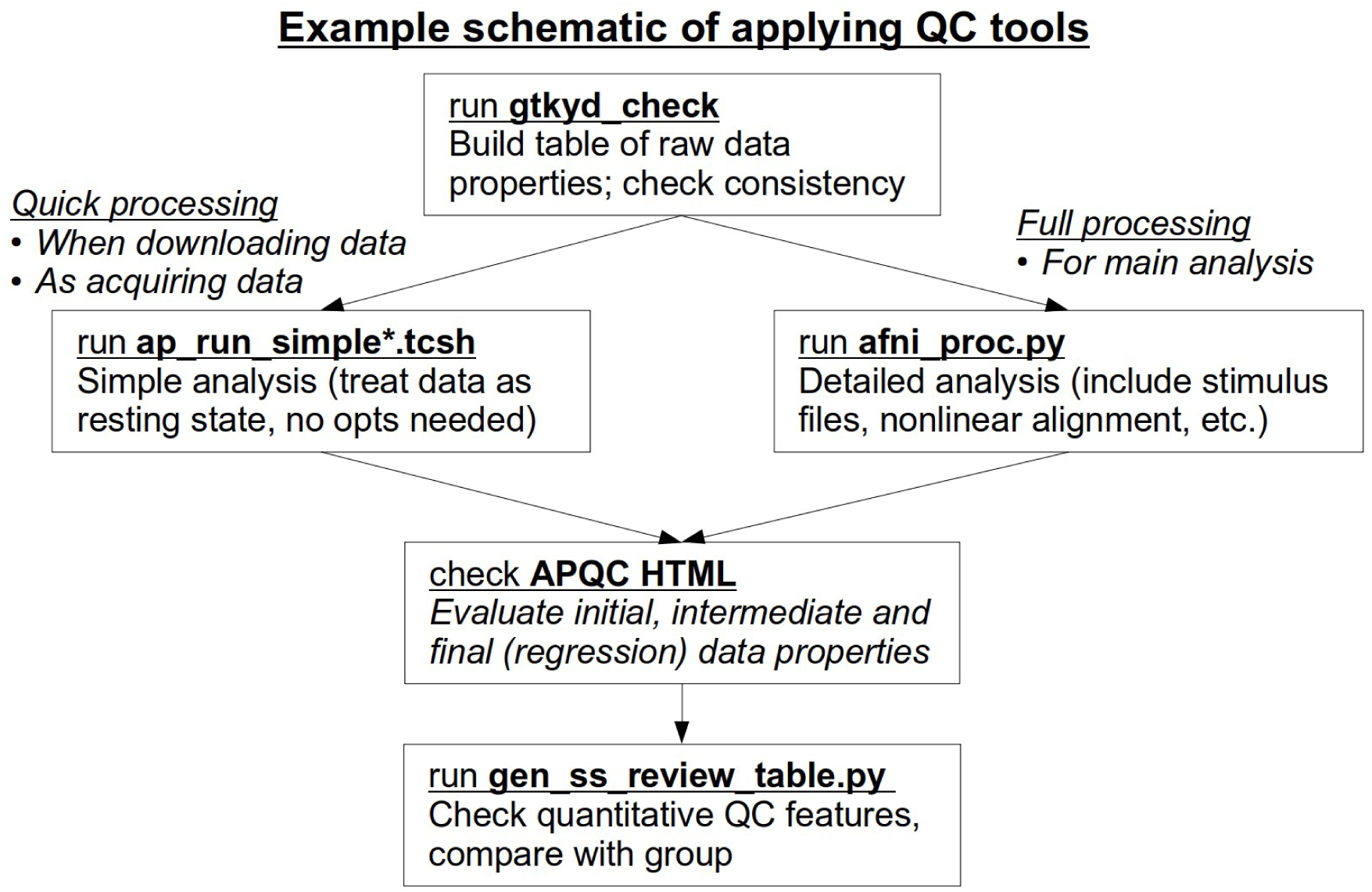
A schematic diagram of ways to integrate major QC tools within AFNI as part of a study. The processing in the second step can be applied in either a detailed sense (full processing, as in preparation for the final study analysis), or in a quick way without needing to specify details (to generate the useful APQC HTML and basic summary quantities in a general manner). Importantly, when acquiring one’s own data, the QC steps should not be left to the end or after large subsets of the data collection have been acquired; it should be applied in a rolling manner, to keep ensuring no artifacts or protocol changes have slipped into the data. Similarly, all downloaded data should be checked for appropriateness—one cannot assume that QC has been done previously, or deeply, or that it evaluated the same properties needed for one’s own study.

Checking the basic data properties is key in all cases, and gtkyd_check provides an efficient way of tabulating data. It is easy to keep rebuilding the table as new datasets are acquired.

To generate the APQC HTML in a quick manner, there are now a pair of "simple" wrapper scripts that run afni_proc.py with essentially no options required from the user. These are called ap_run_simple.tcsh and ap_run_simple_me.tcsh, for processing single- and multi-echo FMRI datasets, respectively. Details are provided in Reynolds et al. (2024). Briefly, the user provides the input subject EPI volumes, and possibly the accompanying T1w anatomical and a reference template name. Then, the scripts process the data like resting state FMRI with default blurring, only affine registration to the template (for speed) and simple motion censoring and regression, which are enough to generate a useful APQC HTML file and accompanying basic quantities. These commands were developed in large part to facilitate QC at the scanner or immediately after acquisition, so the useful HTML summary reports could be made easily. They are also useful for quickly checking properties of a downloaded or shared data collection for fundamental suitability. Of course, one can also run the full afni_proc.py command with detailed choices for a particular study.

Finally, as discussed above, gen_ss_review_table.py can helpfully tabulate and filter the full set of both raw and processing-based properties after running the chosen form of afni_proc.py. The ability to define clear, scriptable criteria for finding subsets of subjects with certain properties is particularly useful. This can also be run repeatedly as new datasets are acquired.

### The need for (more) QC in FMRI research

There are many current discussions in the field about reproducibility of FMRI processing and analysis, such as in a recent editorial "Fostering reproducible fMRI Research" (2017) which states:

> *One means of promoting reproducibility is to ensure transparent reporting of methodological details of study designs, data collection and analytical approaches, as well as any limitations to data interpretaion [sic].*

We strongly agree with this statement and believe that improving quality control—both its practice and communication—has a large role to play in this. As noted above, we believe that the quality control steps are themselves *part* of the processing and not something entirely separate. The described AFNI tools have been developed with this goal in mind. When sharing data, the APQC directories can also be included along with their ratings and comments, as well as with any processing codes, making the QC itself more open. The QC blocks are modular and can be easily described for subject-dropping criteria if necessary, and as demonstrated above these can be integrated directly with programmatic (automated) identification of subjects to exclude. These features serve as one basis of clear and understandable QC reporting.

The availability of public data does not mitigate the need for careful QC. Many open data collections are explicitly uncurated. Even those that may have been curated or checked in *some* way by some other group still require specific evaluations related to any new study. In several projects involving freely available data, some of the present authors have found such problems as: large fractions of subjects having too much motion; slice timing units incorrectly labeled as "seconds" instead of "milliseconds"; overly tight EPI FOVs (truncating the brain); occasional, as well as systematic, left-right flipping between the EPI and T1w volumes; upside-down brains; large amounts of missing or obviously incorrect header information; significant distortions and artifacts; etc. In short, in many cases using data acquired at other centers, by other researchers and with other norms, can require *more* QC oversight by the analyst than if they had acquired it themselves, because the expectations, uses, assumptions, scanner quirks and realtime checks can vary. Even when QC feedback leads collection curators to fix and update the available datasets, researchers will always have varying standards of what are acceptable properties of a data set: for some, censoring more than 10% of EPI time series at a particular threshold is problematic, while for others the fractional cutoff might be 20%, and *others* might use a different motion censor limit; see further discussions in Birn (2023) and Williams et al. (2023). Hence, carrying out QC protocols locally will always be necessary.

One argument often put forward for skipping or reducing QC is that a data collection has too many subjects and QC would take too much time. However, in essentially all cases, the combined amount of time, effort and cost that go into the other aspects of a single subject’s dataset vastly outweighs that of any QC protocol. Consider the subject recruitment process, the scanner time and the actual processing time (each of which is typically more than an hour on its own), as well as a proportion of time used for study planning, grant writing, student training, analysis testing/setup and more. The relative cost of QC itself has only gotten lower as more QC tools have become integrated with processing, such as those with AFNI described here, and is much less than that earlier effort; no one would consider leaving out a processing step, or using linear alignment instead of nonlinear alignment simply because it takes less time*—*why should QC be treated differently?

#### Consider the two scenarios

- If the data and processing have no problems, then browsing the APQC HTML and *gen_ss_review_table.py* output to verify this fact is quite quick and not burdensome;
- If the data and/or processing do have problems that require follow-up, then the need for QC is self-evident and valuable, to avoid biased or misleading results, wasting all the effort of the study.

If the data collection has a large fraction of problematic datasets, then determining that fact saves one from publishing erroneous conclusions, which is a very high cost, indeed. The economics of the scenario always point to performing QC on more datasets in a collection.

Moreover, this points to another important fact: performing deep QC investigations early on in a study is generally necessary to prevent data waste. If there are systematic issues with a scanner setup, protocol or something else, it is better to find out when only 10 subjects have been acquired rather than 100. Furthermore, this helps establish a baseline of properties within the data, and one can also check for any changes in these over time. In all regards, *early and continued QC are important for ensuring valid data collection and processing*.

What to do about large data collections, which can number in the tens of thousands of participants? First, consider that while quality control of this collection might appear a large effort, actually everything about such collections is a large effort, and we must consider the relative amounts. As noted above, compared to grant writing, lab member training, acquisition time and processing time, quality control is only a small fraction of the average work that goes into a subject’s data. If 1 million person-hours have gone into a data collection already, is the fact that adding quality control bumps the number up to 1.01 million person-hours prohibitive? This effort should be weighed against the risk of all 1 million person-hours being wasted by producing biased or artifact-damaged results if no QC is performed (even before accounting for the additional waste and losses for participant and/or clinical populations). There is a strong case for performing full quality control across even large data collections, and the same holds true for studies at all scales.

That being noted, one might also consider that "industrial scale" data collections could benefit from industry-based ideas of maintaining quality. "Deep" quality control could be performed on a given fraction of datasets (say, 10%) spread throughout the entire collection, with "shallow" QC performed on the rest. That is, one could use the QC tools described for full qualitative and quantitative evaluations on subsets of the data at given intervals, to ensure expected properties at those points. Between those intervals, one could use the quantitative measures from processing (the "qsumm" or "basic" quantities, above) and compare those with the same quantities obtained from the deeply QC’ed subsets to check that those are not appreciably changing. As described above, both gtkyd_check and gen_ss_review_table.py are well-suited for this comparison task, scriptable and highly efficient. If changes are observed, then one can perform deeper QC immediately. Having the "simple" afni_proc.py scripts (ap_run_simple*.tcsh) can facilitate this process, but more in-depth afni_proc.py commands could also be set up. Some particular notes about this:

- Pilot data should be acquired before a study starts, and deep QC should be set up using this.
- Deep QC should always be performed at the start of an acquisition.
- If there are known changes within the acquisition (scanner maintenance, change of head coil, software upgrade, change of personnel), then deep QC should be performed on subsequent data.
- If the study is multi-site, such deep- and shallow-QC should be performed at each.
- For multisite studies, it might also be useful to compare dataset QC measures among the sites themselves, if they are to be merged later.

In this way, the initial deep QC sets a baseline expectation with the datasets, creating a "training set" range of meaningful quantities for the study goals. Shallow QC is used to more cheaply check that a large number of datasets remain within those quantitative ranges. Periodic deep QC is necessary to evaluate and verify the consistency of data properties. The higher the fraction of deep QC, the more confident researchers can be. Even for a single site study, involving multiple people in QC discussions and evaluations is helpful, which requires building consensus. In practice, this should improve the long-term maintenance and stability of QC, such as if some evaluators leave and new ones join.

Several existing projects with very large data collection size have described performing combinations of manual and automated QC integrated with their acquisition protocols. The UK Biobank description focused on using machine learning and quantitative metrics to perform primarily automated quality control for FMRI (Alfaro-Almagro et al., 2018). In contrast, Hagler et al. (2019) described a combination of automated quantitative summary parameters and manual QC, such as using montages of images, in the ABCD study. Monereo-Sanchez (2021) described a comparison of QC procedures for a relatively large sampling of structural datasets from the Maastricht Study, concluding that those including manual/visual QC resulted in the least unexplained variance; however, they also noted this required training to reduce bias or subjectivity (and for related comments on FMRI specifically, see Williams et al., 2023). Ma et al. (2022) applied outlier detection to both the multimodal UK Biobank and Human Connectome Project data collections, followed by visual assessment of those extrema. While this project was focused on finding rare imaging phenotypes within large cohorts, it also performed combined quantitative and qualitative QC evaluations in its two-step approach (and noted particularly low test-retest reliability in their resting state FMRI data parameterization, attributed to subject motion and other influences). In all cases, when reusing the data for separate projects, it would still be advisable to perform one’s own evaluation for their suitability to the new application.

### QC tools across the field

There are many QC tools besides those in AFNI within the field of neuroimaging. Examples of FMRI QC tools in other softwares and pipelines include: the Conn toolbox (Whitfield-Gabrieli and Nieto-Castanon, 2012), C-PAC (Craddock et al., 2013), DPABISurf (Yan et al., 2019), DPARSF (Yan et al., 2010), ENIGMA HALFpipe (Waller et al., 2022), fMRIPrep (Esteban et al., 2019), FreeSurfer (Fischl and Dale, 2000), FSL (Smith et al., 2004), FSLeyes (McCarthy, 2012), LONI QC pipeline (Rex et al. 2003), MRIQC (Esteban et al., 2017), pyfMRIqc (Williams and Lindner 2020), SPM12 (https://www.fil.ion.ucl.ac.uk/spm/; Ashburner et al., 2012), and SynthStrip (Hoopes et al., 2022). Many of these have overlapping features (including the fact that some tools implement other software packages under the hood), as observed in the FMRI Open QC Project (see Taylor et al., 2023a). For example, MRIQC also allows users to make ratings and save them in its HTML document. But each tool also has its own emphases, ranges/subsets of processing coverage, implementations and philosophies.

There are too many aspects to compare across software packages and projects. We refer people instead to the documentation of the individual packages, as well as to the useful examples in the demonstration papers for those participated in the FMRI Open QC Project: AFNI (Birn, 2023); SPM (Di and Biswal, 2023); R scripts and AFNI (Etzel, 2023); AFNI (Lepping et al., 2023); DPABISurf, DPARF and fMRIPrep (Lu and Yan, 2023); CONN, SPM12 and FSLeyes (Morfini et al., 2023); MRIQC and fMRIPrep (Provins et al., 2023); AFNI (Reynolds et al., 2023); AFNI (Teves et al., 2023); and pyfMRIqc (Williams et al., 2023). These provide useful QC tutorials on many of these packages, with lists of specific QC items, detailed descriptions, and example cases.

### Design choices in the AFNI QC features

The QC features presented here have been constructed with the goal of understanding the properties of the input data as fully as possible. These have been developed over time with a primary focus on visualizing data in each of its raw, intermediate and final stages, evaluating many features and processing steps (e.g., alignment, signal patterns, local tissue contrast, etc.). Quantitative checks have also been integrated with this, building on the qualitative understanding and also facilitating it. Because the utilized measures have been developed in conjunction with both active pipeline development and the experience of qualitative assessment, they have been curated to have meaningful relations to the data and elucidate key features. Most features directly target known problems that have been observed in data, particularly those in the "warns" block of the APQC HTML (collinearity in models, left-right inconsistency, anomalous variance lines, and more), as well as helpfully summarize useful properties (average values within masks, GCOR, degree of freedom counts, etc.).

In addition to (and largely due to) their developmental ties, the qualitative and quantitative assessment features reinforce and synergize each other in the QC evaluations. FMRI datasets are acquired with an intricate array of physics and are themselves noisy and complicated; there are also a wide variety of processing steps applied to them before they can be interpreted or used in further analyses. To date, evaluating the datasets from start to finish, typically with a combination of qualitative and quantitative criteria at each stage, remains the best way to reduce both the effects of artifacts and the chances of undetected problems affecting final results. The goal of QC development is then to provide a useful set of checks and tools for understanding the data in a manner that balances efficiency with completeness. The APQC HTML aims for this by providing a systematic set of QC blocks, images and quantities within a single interface. It is worth noting that these checks cover the gamut of raw data through linear regression modeling, providing a relatively comprehensive overview of complete single subject processing.

With the recent addition of user interactive features (AV, NV, IC, GV and QC rating/comment buttons), users can dive into deeper investigations relatively quickly, to help resolve questions that arise. Being able to save comments and ratings during QC evaluation also benefits the efficiency of the process and the ability to share and discuss findings further. The gen_ss_review_table.py functionality facilitates tabulating and comparing results across a group efficiently, as well as programmatically identifying subjects with particular properties.

Another important addition has been the tables of local properties of ROI shape and TSNR. As stressed in Reynolds et al. (2023), the evaluation of the suitability of data is closely tied to an explicit study’s goals. Researchers must ensure that they have the desired coverage and spatial resolution to study their anatomical regions of interest. Parcellations are defined in many ways, and researchers might find that a given dataset is only suitable to particular granularity. The default MNI APQC atlas was intentionally created using parcellations of differing fineness: the Glasser HCP (360 regions; Glasser et al., 2016) and Julich Brain atlas (Amunts et al., 2020); and the larger, functionally-defined Schaefer-Yeo atlas (100 regions from 17 networks; Schaefer et al., 2018), specifically the AFNI-refined ROIs from (Glen et al. 2021). When acquiring one’s own data, we expect that this table can usefully inform the tuning of acquisition protocol, to ensure that spatial resolution is actually high enough to resolve one’s desired ROIs (akin to meeting a Nyquist criterion of sampling in time series analysis).

Similarly, FMRI signal strength is not constant across the brain, and researchers must be careful to have usable data in their regions of interest. There are several brain areas known to be particularly susceptible to signal distortion, dropout and attenuation, such as the ventromedial prefrontal cortex (VMPFC), subcortex and temporal lobes. Knowing the TSNR properties of these or any other key regions is important for evaluating data for a study. Having either low overall magnitudes or strong changes/gradients in TSNR would suggest data quality problems in those locations, which might make results using those unreliable. These tables in the APQC HTML should greatly facilitate checks of public data for given research projects. More fundamentally, we hope they are implemented in pilot studies to evaluate the data at the acquisition stage, feeding back into improving the protocol itself wherever possible.

The number, type and content of QC blocks will surely grow over time. For example, researchers can use the "surf" block in *afni_proc.py* to perform surface-based FMRI processing with SUMA (Saad et al., 2004); the scriptable images from SUMA can also be incorporated into the APQC HTML. Additionally, *afni_proc.py* also can include processing associated with B0 inhomogeneity distortion correction, including fieldmap correction (Jezzard and Balaban, 1995) and blip-reversed unwarping (Chang and Fitzpatrick, 1992; Andersson et al., 2003); in each case, the quality of distortion correction can be shown in a manner similar to the comparisons of the present "ve2a" and "va2t" alignment blocks. As noted in the Methods, QC information from other software that has been integrated into *afni_proc.py* has already been included in the APQC HTML (the tedana MEICA calculation, in the "mecho" QC block); the addition of other software dependencies would likely lead to further QC additions in a similar way. In general, as with most processing tools, one can only expect functionality to evolve with continued user-feedback and collaboration.

### Limitations and future work

The processing framework for *afni_proc.py* can set up a wide variety of processing pipelines, including those that include surface-based analysis (see Reynolds et al., 2024 for an example). At present, the APQC HTML does not contain systematic images of the surface-based parts of the processing. Work is underway to include this, through adding automatic snapshot features to SUMA (Saad et al., 2004) as well as integration with NiiVue-based visualization within the HTML.

FMRI acquisition and study paradigms continue to grow and change, and recently the FMRI Open QC project has spurred QC discussions across the neuroimaging community for sharing more QC suggestions (see Taylor et al., 2023a, and op cit). We have no doubt that further QC features will be integrated with those already created in AFNI and with afni_proc.py. We plan to further integrate cross-subject properties across data collections. We aim in particular to extend important visualization and qualitative understanding to improve development of more quantitative metrics to guide researchers’ understanding of data.

## CONCLUSIONS

We have described several layers of FMRI data and processing quality control within the AFNI software package. Viewing "quality control" as the broad project of understanding the data, many different features need to be evaluated, both qualitative and quantitative. AFNI’s primary FMRI pipeline tool, *afni_proc.py*, contains several layers of QC features, including clear provenance of processing (for clarity of what is being examined), retention of intermediate data (to be able to investigate in depth), multiple QC scripts (guiding important checks), text files of automatic checks (e.g., regression matrix warnings), a quantitative summary report (including both input and derived quantities), and a systematic HTML report for detailed but efficient investigation (combining both qualitative and quantitative information). Having all of the QC blocks in a single place for a given subject makes it easy to quickly understand the "story" of the processing, and to identify potential problems by jumping between different pieces of information; this has further been integrated with being able to choose any particular QC item and navigate efficiently across subjects evaluating it. We described helpful new functionality that facilitates user interaction, for even more in-depth investigations in an efficient manner, and saving of QC ratings and notes. These quantitative and qualitative outputs can be combined into a single summary table with gen_ss_review_table.py, which can also apply "exclusion criteria" programmatically. We believe that these tools will greatly facilitate the understanding and communication of FMRI data quality evaluations.

## ACKNOWLEDGMENTS

PAT, RCR, DRG, JKR and RWC are supported by the NIMH Intramural Research Programs of the NIH, HHS, USA (ZICMH002888), as is DMN (ZICMH002968). CR is supported by NIH RF1-MH133701. TH is supported by the Wellcome Centre for Integrative Neuroimaging (Oxford, UK) via funding from the Wellcome Trust (203139/Z/16/Z and 203139/A/16/Z). This work utilized the computational resources of the NIH HPC Biowulf cluster (http://hpc.nih.gov).

## ETHICS

The datasets used in this study are publicly available and were not collected by the investigators.

## DATA AND CODE AVAILABILITY

The data are available on OSF (https://osf.io/un56d/), and the processing commands are available on GitHub (https://github.com/afni/apaper_apqc_etal).

## AUTHOR CONTRIBUTIONS

PT: conceptualization, formal analysis, methodology, software, visualization, writing – original draft preparation; DRG: methodology, software, visualization, writing – review and editing; GC: methodology, software, visualization, writing – review and editing; RWC: methodology, software, visualization, writing – review and editing; DMN: methodology, visualization, writing – review and editing; CR: software, visualization, writing – review and editing; TH: software, visualization, writing – review and editing; JKR: methodology, software, visualization, writing – review and editing; RCR: conceptualization, formal analysis, methodology, software, visualization, writing – original draft preparation;

## CONSENT FOR PUBLICATION

All authors consent to publication.

## Footnotes

1 The data are available on OSF (https://osf.io/un56d/), and the processing commands are available on GitHub (https://github.com/afni/apaper_apqc_etal).

2 Here and below, we use the shell syntax of ${X} to denote some variable "X", with the variable’s name suggesting what it represents. For example, ${subj} represents the subject ID.

3 If the APQC HTML was opened without the Flask modules available, whether via the open_apqc.py command, another command line tool or simply clicking on the file, the page information is viewable, but previous QC ratings and comments will not be visible, nor will new items be stored. The "RATE" text in these cases will be gray and struck out with a red line.

4 Enorm is analogous to the "FD" parameter used in other software. The main difference is that the former is an *L^2^*- norm of motion parameters, while the latter is an *L^1^*-norm. The *L^2^*-norm is preferred in AFNI because it has a closer formulation to how distances are actually calculated and is also more sensitive to large changes in any individual motion parameter.

5 For more discussion of this, particularly with regard to processing choices, see Appendix A of Reynolds et al. (2024).

6 https://afni.nimh.nih.gov/pub/dist/doc/htmldoc/codex/main_toc.html

7 See for example, @Install_APMULTI_Demo1_rest and demos listed here: https://afni.nimh.nih.gov/pub/dist/doc/htmldoc/nonhuman/main_toc.html

8 See https://www.youtube.com/c/afnibootcamp, https://afni.nimh.nih.gov/pub/dist/doc/htmldoc/educational/main_toc.html

